# LUBAC assembles a signaling platform at mitochondria for signal amplification and shuttling of NF-ĸB to the nucleus

**DOI:** 10.1101/2022.05.27.493704

**Authors:** Zhixiao Wu, Lena A. Berlemann, Verian Bader, Dominik Sehr, Eva Eilers, Alberto Covallero, Jens Meschede, Lena Angersbach, Cathrin Showkat, Jonas B. Michaelis, Christian Münch, Bettina Rieger, Dmitry Namgaladze, Maria Georgina Herrera, Fabienne C. Fiesel, Wolfdieter Springer, Marta Mendes, Jennifer Stepien, Katalin Barkovits, Katrin Marcus, Albert Sickmann, Gunnar Dittmar, Karin B. Busch, Dietmar Riedel, Marisa Brini, Jörg Tatzelt, Tito Cali, Konstanze F. Winklhofer

## Abstract

Mitochondria are increasingly recognized as cellular hubs to orchestrate signaling pathways that regulate metabolism, redox homeostasis, and cell fate decisions. Recent research revealed a role of mitochondria also in innate immune signaling, however, the mechanisms of how mitochondria affect signal transduction are poorly understood. Here we show that the NF-ĸB pathway activated by TNF employs mitochondria as a platform for signal amplification and shuttling of activated NF-ĸB to the nucleus. TNF induces the recruitment of HOIP, the catalytic component of the linear ubiquitin chain assembly complex (LUBAC), and its substrate NEMO to the outer mitochondrial membrane, where M1- and K63-linked ubiquitin chains are generated. NF-ĸB is locally activated and transported to the nucleus by mitochondria, resulting in an increase in mitochondria-nucleus contact sites in a HOIP-dependent manner. Notably, TNF-induced stabilization of the mitochondrial kinase PINK1 contributes to signal amplification by antagonizing the M1-ubiquitin-specific deubiquitinase OTULIN.

## INTRODUCTION

In addition to their bioenergetic and biosynthetic functions, mitochondria have emerged as potent and versatile signaling organelles to adapt metabolism to cellular demands, to communicate mitochondrial fitness to other cellular compartments, and to regulate cell death and viability. Increasing evidence implicates mitochondria in innate immune signaling, illustrating that mitochondria have been co-opted by eukaryotic cells to react to cellular damage and to promote efficient immune responses. For example, invading pathogens expose pathogen-associated molecular patterns (PAMPs), such as viral RNA, which is recognized by cytoplasmic RLRs (retinoic acid inducible gene-I-like receptors). Binding of viral RNA to RLRs induces a conformational change, allowing their recruitment to the mitochondrial RLR adaptor protein MAVS (mitochondrial antiviral signaling protein). Subsequent multimerization of MAVS at mitochondria results in the activation of the transcription factors and IRF (interferon-regulatory factor) 3, IRF7, and NF-ĸB (nuclear factor kappa B) that regulate the expression of type 1 interferons and pro-inflammatory cytokines (Ablasser and Hur, 2020; Chowdhury et al., 2022; Rehwinkel and Gack, 2020; Seth et al., 2005).

NF-κB is activated in several innate immune paradigms and is well known for its role in pathological conditions, where prolonged or profuse NF-κB activation in reactive immune or glial cells causes inflammation. However, NF-κB has important physiological functions in most -if not all-cell types. In the nervous system, NF-κB maintains neuronal viability and regulates synaptic plasticity (Dresselhaus et al., 2018; Meffert et al., 2003; Neidl et al., 2016; O’Riordan et al., 2006; Salles et al., 2015). Constitutive NF-κB activation by para-or autocrine loops is required for neuronal survival, and stress-induced transient NF-κB activation confers protection from various neurotoxic insults (Bhakar et al., 2002; Blondeau et al., 2001). Accordingly, TNF has been identified as a neuroprotective cytokine that prevents neuronal damage under various stress conditions (Cheng et al., 1994; Turrin and Rivest, 2006).

Although TNF can induce apoptotic or necroptotic cell death in certain pathological conditions, it usually protects from cell death. Engagement of TNF with its receptor TNFR1 at the plasma membrane results in the formation of signaling complex I that activates the transcription factor NF-ĸB (typically p65/p50 heterodimers) in a tightly regulated manner, involving multiple phosphorylation and ubiquitination events (Annibaldi and Meier, 2018; Brenner et al., 2015; Hayden and Ghosh, 2014; Varfolomeev and Vucic, 2018). A crucial step in the activation process is the ubiquitination of NEMO (NF-ĸB essential modulator) by HOIP, the catalytic component of the linear ubiquitin chain assembly complex (LUBAC), and the binding of NEMO to linear ubiquitin chains by its UBAN (ubiquitin-binding domain in ABIN and NEMO) domain (Haas et al., 2009; Rahighi et al., 2009; Tokunaga et al., 2009). These events induce a conformational change and oligomerization of NEMO, activating the associated kinases IKKα and IKKβ. IKKα/β phosphorylate the NF-ĸB inhibitor IĸBα, which is subsequently modified by K48-linked ubiquitin chains and degraded by the proteasome. NF-ĸB heterodimers, liberated from their inhibitory binding to IĸBα, are then translocated from the cytoplasm to the nucleus, where they increase the expression of anti-apoptotic proteins or pro-inflammatory cytokines in cell type-and context-specific manner.

We have previously shown that the E3 ubiquitin ligase Parkin depends on this pathway to protect from stress-induced neuronal cell death (Henn et al., 2007; Muller-Rischart et al., 2013). Parkin modifies NEMO with mono-ubiquitin or K63-linked ubiquitin chains and facilitates subsequent M1-linked ubiquitination of NEMO by HOIP, supporting the concept that linear ubiquitin chains are preferentially assembled as heterotypic K63/M1-linked chains in innate immune signaling pathways (Cohen and Strickson, 2017; Emmerich et al., 2016; Emmerich et al., 2013; Fiil et al., 2013; Hrdinka et al., 2016). Notably, in the absence of either HOIP or NEMO, Parkin cannot prevent stress-induced cell death (Muller-Rischart et al., 2013).

Linear or M1-linked ubiquitination is characterized by the head-to-tail linkage of ubiquitin molecules via the C-terminal carboxyl group of the donor ubiquitin and the N-terminal methionine of the acceptor ubiquitin (Dittmar and Winklhofer, 2019; Fiil and Gyrd-Hansen, 2021; Fuseya and Iwai, 2021; Jahan et al., 2021; Oikawa et al., 2020; Spit et al., 2019; Tokunaga and Ikeda, 2022). Formation and disassembly of linear ubiquitin chains are highly specific processes. HOIP is the only E3 ubiquitin ligase that generates linear ubiquitin chains, whereas OTULIN is the only deubiquitinase that exclusively hydrolyses linear ubiquitin chains (Lork et al., 2017; Verboom et al., 2021; Weinelt and van Wijk, 2021). Thus, M1-linked ubiquitination can be studied with unique specificity, in contrast to other types of ubiquitination. Mass spectrometry data on TNF-induced signaling networks and the interactive profile of HOIP and OTULIN directed our interest towards mitochondria (Kupka et al., 2016a; Stangl et al., 2019; Wagner et al., 2016). Here we show that TNF induces remodeling of the outer mitochondrial membrane by assembling an NF-ĸB signaling platform that facilitates nuclear translocation of activated NF-ĸB. HOIP plays a crucial role in this process by generating M1-linked ubiquitin chains and stabilizing the mitochondrial kinase PINK1, which contributes to NF-ĸB signal amplification.

## RESULTS

### TNF induces recruitment of LUBAC and the assembly of linear ubiquitin chains at mitochondria

To get insight into the subcellular localization of OTULIN and its interactive profile, we performed co-immunoprecipitation of HA-tagged OTULIN expressed in HEK293T cells coupled to mass spectrometry. Strict filtering criteria were applied in two independent experiments, including only proteins, which were detected by at least two unique peptides that were not present in the negative controls. By this approach, 59 potential OTULIN-interacting proteins were identified (Suppl. Fig. 1A). An analysis of the subcellular localization of OTULIN-interacting proteins indicated several mitochondrial proteins (Suppl. Fig. 1B), consistent with interactors screens previously performed with OTULIN and HOIP (Kupka et al., 2016a; Stangl et al., 2019). Moreover, mitochondrial proteins were found in a study on proteome-wide dynamics of phosphorylation and ubiquitination in TNF-induced signaling (Wagner et al., 2016). These findings prompted us to explore a possible mitochondrial localization of OTULIN and HOIP. Mitochondria from HEK293T cells were isolated and purified by subcellular fractionation using differential centrifugation followed by isopycnic density gradient centrifugation. Immunoblotting of purified mitochondrial fractions revealed that both OTULIN and HOIP co-purified with mitochondria (Fig. 1A). We wondered whether OTULIN antagonizes M1-ubiquitination at mitochondria and therefore silenced OTULIN expression by RNA interference. Immunoblotting of mitochondrial fractions using the M1-ubiquitin-specific antibodies 1E3 or 1F11/3F5/Y102L (Matsumoto et al., 2012) revealed a strong M1-ubiquitin-positive signal at mitochondria isolated from OTULIN-silenced cells, indicating that OTULIN regulates mitochondrial M1-ubiquitination (Fig. 1A). Based on the fact that TNF activates LUBAC (Haas et al., 2009; Rahighi et al., 2009; Tokunaga et al., 2009) and the identification of mitochondrial proteins in the proteome analysis of TNF signaling (Wagner et al., 2016), we were wondering if TNF induces linear ubiquitin chain formation at mitochondria. M1-linked ubiquitin chains were detected 15 min after TNF treatment at mitochondria isolated from various cells, such as HEK293T, HeLa, SH-SY5Y cells, and mouse embryonic fibroblasts (MEFs) (Fig. 1A, B). TNF-induced mitochondrial M1-ubiquitination was not observed in HOIP CRISPR/Cas9 knockout (KO) cells, confirming HOIP-dependent linear ubiquitination (Fig. 1C). Co-localization of M1-linked ubiquitin with mitochondria after TNF treatment was also seen by immunocytochemistry followed by super-resolution structured illumination microscopy (SR-SIM) (Fig. 1D). Consistent with an increase in mitochondrial M1-ubiquitination, the abundance of all three LUBAC components, HOIP, HOIL-1L and SHARPIN, was increased at mitochondria after TNF treatment (Fig. 1E). Immunoblotting of mitochondrial fractions showed that M1-ubiquitination at mitochondria is a fast and transient response with maximum intensity after 15 min that is paralleled by an increase in K63-ubiquitination (Fig. 1F). This is in line with the observation that linear ubiquitin chains are preferentially assembled as heterotypic K63/M1-linked chains (Cohen and Strickson, 2017; Emmerich et al., 2016; Emmerich et al., 2013; Fiil et al., 2013; Hrdinka et al., 2016).

**Figure 1.**
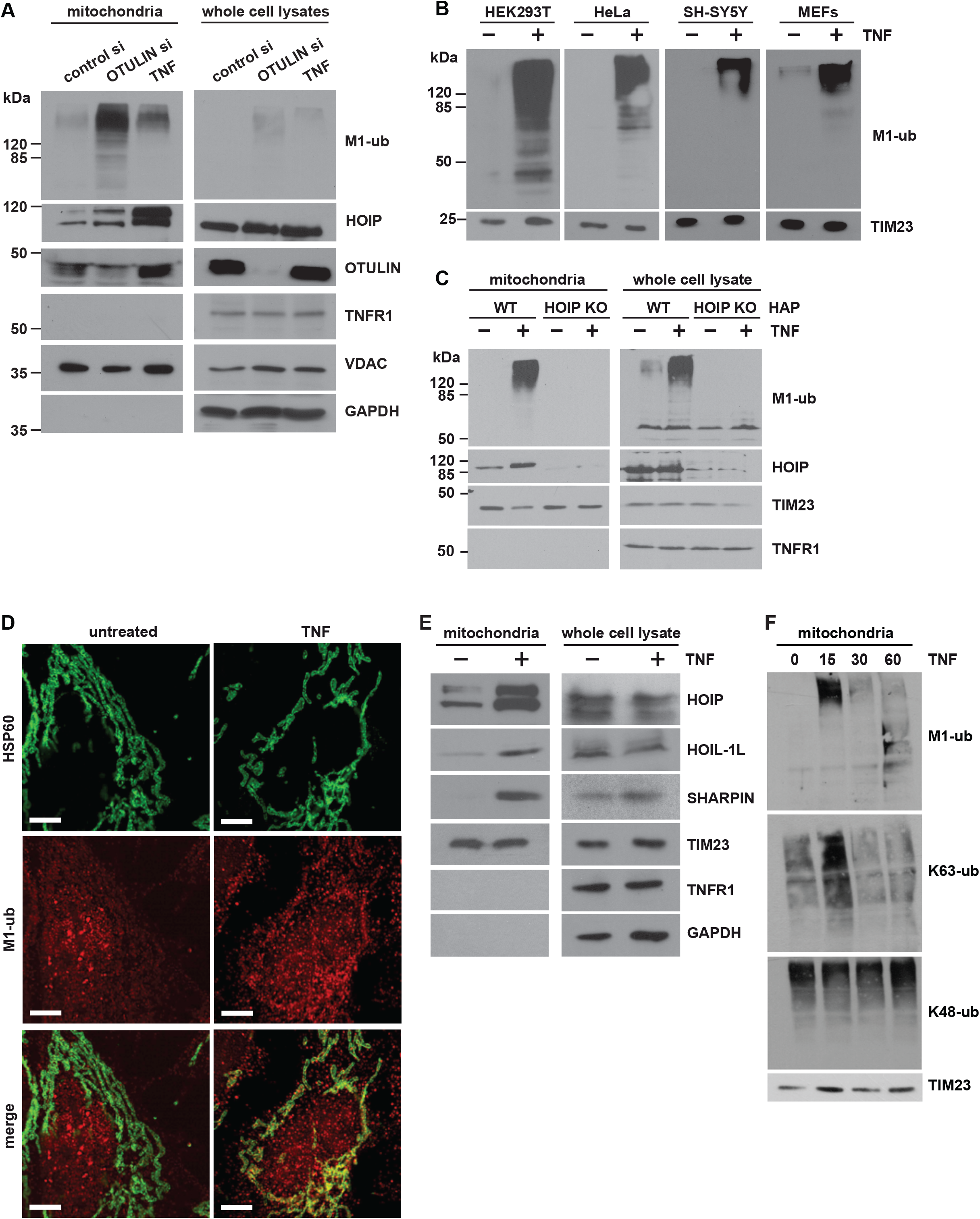
TNF induces the formation of M1-linked ubiquitination at mitochondria. **A. A fraction of OTULIN is localized at mitochondria where it suppresses M1-linked ubiquitination**. HEK293T cells were transfected with control or OTULIN siRNA. 72 h after transfection or 15 min after TNF treatment (25 ng/ml) the cells were harvested. Mitochondria were isolated by differential centrifugation and purified by ultracentrifugation using an OptiPrep™ density gradient. Mitochondrial fractions (38% of total cell lysate) and whole cell lysates (2%) were analyzed by immunoblotting using the antibodies indicated. **B. TNF induces M1-ubiquitination at mitochondria in various cell types**. The indicated cell types were treated with TNF (25 ng/ml, 15 min) and purified mitochondrial fractions analyzed as described in A. **C. TNF-induced mitochondrial M1-ubiquitination does not occur in HOIP-deficient cells**. Wildtype and HOIP-KO HAP cells were treated with TNF and analyzed as described in A. **D. Mitochondria and M1-linked ubiquitin co-localize after TNF treatment**. SH-SY5Y cell were treated with TNF (25 ng/ml, 15min), fixed, stained with antibodies against HSP60 (green) and M1-ubiquitin (red) and analyzed by SR-SIM. Scale bar, 5 µm. **E. TNF induces recruitment of LUBAC components to mitochondria**. HEK293T cells were treated with TNF (25 ng/ml, 15 min) and analyzed as described in A. **F. TNF induces a fast and transient increase in M1- and K63-specific ubiquitination at mitochondria**. HEK 293T cells were treated with TNF (25 ng/ml) for the indicated time and the mitochondrial fractions were analyzed by immunoblotting using M1-, K63-, and K48-specific ubiquitin antibodies.

### TNF stabilizes PINK1 at mitochondria

Next, we addressed the functional consequences of TNF-induced mitochondrial ubiquitination. Ubiquitination of mitochondrial outer membrane proteins has been linked to a mitochondrial quality control pathway, resulting in the clearance of mitochondria by mitophagy (Goodall et al., 2022; Harper et al., 2018; Montava-Garriga and Ganley, 2020; Nguyen et al., 2016; Pickles et al., 2018; Whitworth and Pallanck, 2017). In this pathway, depolarization of the inner mitochondrial membrane leads to the stabilization of the mitochondrial kinase PINK1 at the outer mitochondrial membrane. PINK1 phosphorylates ubiquitin and the ubiquitin-like domain of the E3 ubiquitin ligase Parkin at serine 65, thereby activating Parkin. Parkin-mediated ubiquitination of outer membrane proteins then recruits the autophagic machinery to eliminate damaged mitochondria. When the mitochondrial membrane potential is intact, full-length PINK1 is imported by the translocases of the outer and inner mitochondrial membranes, the TOM and TIM23 complexes, and then PINK1 is processed to a 60 kDa intermediate species by the mitochondrial processing peptidase (MPP) in the matrix. Subsequently, the presenilin-associated rhomboid-like protein (PARL) in the inner membrane generates a 52 kDa mature PINK1 fragment (Deas et al., 2011; Greene et al., 2012; Jin et al., 2010; Meissner et al., 2011; Shi et al., 2011; Whitworth et al., 2008). Proteolytically processed PINK1 is retro-translocated to the mitochondrial outer membrane, where it is ubiquitinated and rapidly degraded by the proteasome (Liu et al., 2017; Yamano and Youle, 2013). Whereas non-imported full-length PINK1 has been linked to mitophagy induction, little is known about the function of mature PINK1 and the conditions that increase its abundance.

The artificial tethering of unbranched linear ubiquitin chains to the mitochondrial outer membrane was shown to induce mitophagy (Yamano et al., 2020), we therefore tested whether mitochondrial M1-ubiquitination induced by TNF has an impact on mitophagy. We first compared the relative amounts of PINK1 species in untreated, TNF-stimulated and CCCP (carbonyl cyanide 3-chlorophenylhydrazone)-treated cells. As expected, dissipation of the proton-motive force across the mitochondrial inner membrane by CCCP (10 µM, 90 min) stabilized the unprocessed 63 kDa PINK1 species due to an impeded mitochondrial import. Interestingly, TNF stabilized both the unprocessed and processed PINK1 species after 15 min (Fig. 2A). A kinetic analysis of TMRE (tetramethylrhodamine ethyl ester) fluorescence in TNF-treated cells indicated a mild transient increase in the membrane potential between 15 and 60 min (Fig. 2B), revealing that PINK1 in TNF-treated cells was not stabilized by a dissipation of the mitochondrial membrane potential. Consistent with this result, TNF-induced PINK1 stabilization at mitochondria did not induce mitophagy. Quantitative assessment of mitophagy by flow cytometry using the fluorescent reporter mt-Keima did not reveal changes in mitophagic activity after 30 min and 16 h of TNF treatment (Fig. 2C).

**Figure 2.**
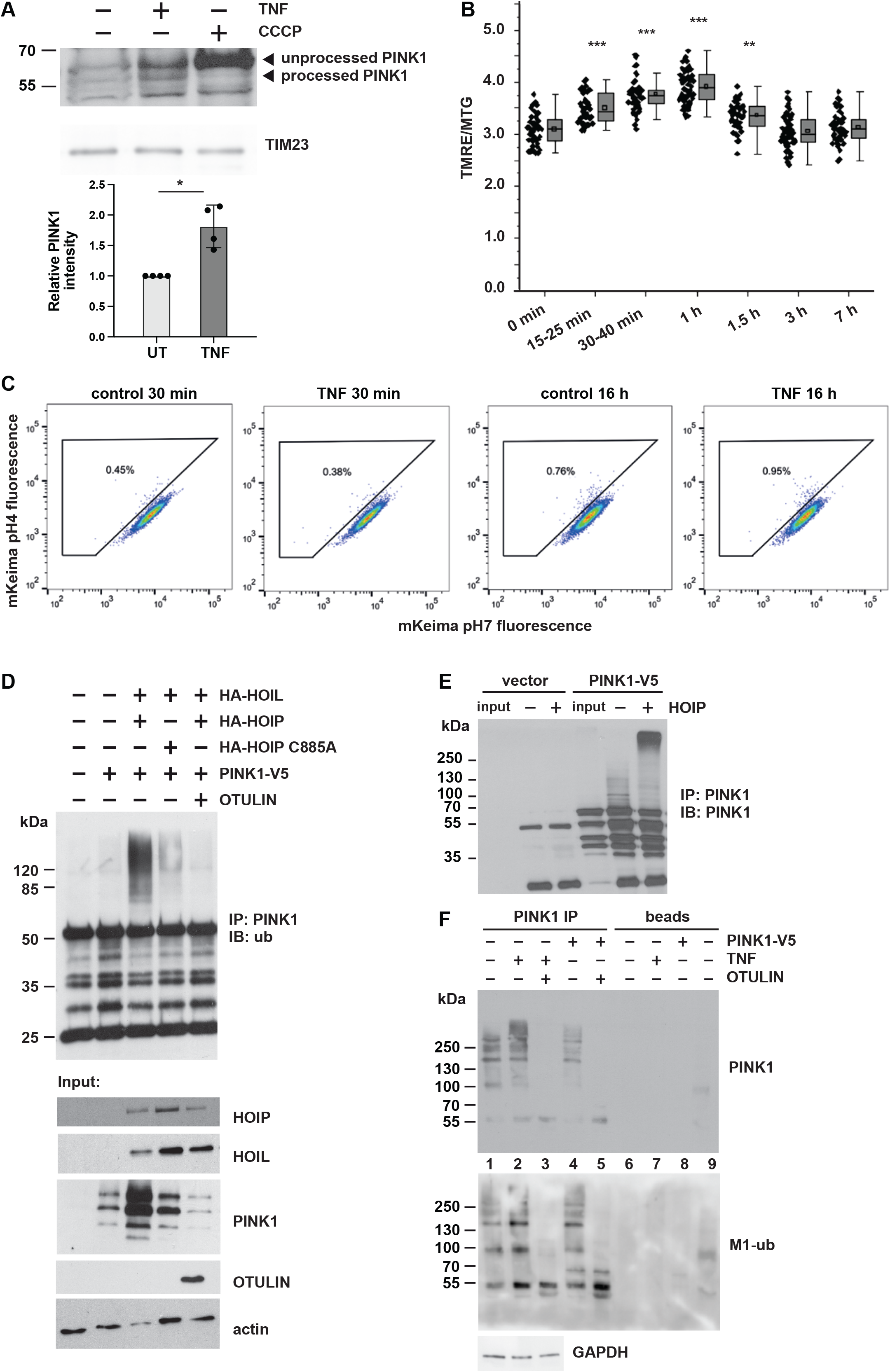
PINK1 is stabilized at mitochondria by LUBAC-mediated ubiquitination. **A. PINK1 is stabilized by TNF treatment**. HEK293T cells were treated with TNF (25 ng/ml, 15 min) or CCCP (10 µM, 90 min) before harvesting. Purified mitochondrial fractions were analyzed by immunoblotting using the indicated antibodies (upper panel). Quantification of PINK1-specific signals normalized to TIM23 signals (lower panel). Data represent the mean values with standard deviations of 4 independent experiments. *p < 0.05. A two-tailed non-parametric Mann-Whitney U-test was used to analyze statistical significance. **B. TNF-induced PINK1 stabilization is not associated with a decrease in the mitochondrial membrane potential**. HeLa cells were treated with TNF (25 ng/ml, 15 min) and the mitochondrial membrane potential was assessed by the fluorescence signal intensity ratio of TMRE and MitoTracker™ Green (MTG), n=2. **C. TNF-induced PINK1 stabilization does not induce mitophagy**. HeLa cells expressing mt-mKeima were treated with TNF (25 ng/ml, 15min) for 30 min or 16 h. Analysis was done by flow cytometry gating of lysosomal and neutral mt-mKeima. **D. Catalytically active HOIP ubiquitinates overexpressed PINK1**. HEK293T cells were transfected with the plasmids indicated. One day later, the cells were harvested under denaturing conditions and PINK1 was immunoprecipitated via the V5 tag followed by immunoblotting using ubiquitin antibodies. **E. Recombinant HOIP ubiquitinates PINK1 *in vitro***. PINK1 immunoprecipitated from transiently transfected HEK293T cells via the V5 tag was incubated with recombinant mouse Ube1, UBE2L3, C-terminal HOIP and ubiquitin for *in vitro* ubiquitination. The samples were then analyzed by immunoblotting using V5 antibodies. **F. Endogenous PINK1 is modified with M1 ubiquitin chains after TNF treatment**. HEK293T cells were treated with TNF (25 ng/ml, 15min), then endogenous PINK1 was immunoprecipitated from mitochondrial fractions. We also included cells mildly overexpressing PINK1-V5 (lanes 4, 5, 8), to make sure that the immunoreactive bands seen for endogenous PINK1 indeed correspond to PINK1. As a control for the presence of M1-linked ubiquitin chains, immunoprecipitated PINK1 was treated with recombinant OTULIN. As controls for the specificity of the immunoprecipitation, beads only (lanes 6 – 8) and beads plus IgG (lane 9) were included. The samples were analyzed by immunoblotting using PINK1 and M1-ubiquitin antibodies. For the immunoprecipitation of overexpressed PINK1, only half the amount of cells were used in comparison to the immunoprecipitation of endogenous PINK1.

It has been reported previously that the mature 52 kDa PINK1 species is stabilized by K63-linked ubiquitination induced by NF-κB pathway activation (Lim et al., 2015). We therefore wondered whether PINK1 is a target for linear ubiquitination. Co-immunoprecipitation experiments indicated that PINK1 interacts with HOIP (Suppl. Fig. 2A, B). To map the domains of HOIP mediating this interaction, we employed transiently transfected cells. V5-tagged PINK1 co-purified with full-length HOIP and the N-terminal part of HOIP (aa 1-697), but not with the C-terminal part of HOIP (aa 697-1072) (Suppl. Fig. 2A, B). We then tested different HOIP constructs lacking N-terminal domains to more precisely define the region of HOIP that interacts with PINK1. PINK1 co-immunoprecipitated with the 1-697 aa HOIP construct and to a lesser extent with the 1-475 aa construct, which lacks the UBA domain, suggesting that the UBA domain directly or indirectly contributes to the interaction of HOIP with PINK1 (Suppl. Fig. 2C, D). Of note, immunoblotting of cellular lysates indicated that PINK1 abundance was increased in the presence of either full-length HOIP or N-terminal HOIP (1-697 aa), suggesting that complex formation contributes to PINK1 stabilization (Suppl. Fig. 2B, D, input). In support of PINK1 interacting with endogenous HOIP, both proteins were detected in the 450 - 720 kDa fractions (corresponding to the size of LUBAC) upon size exclusion chromatography of cellular lysates (Suppl. Fig. 2E). In a next step, PINK1 ubiquitination was analyzed in cells overexpressing HOIP and HOIL-1 to increase LUBAC activity. The cells were harvested under denaturing conditions and then PINK1 was affinity-purified by its V5 tag and subjected to immunoblotting using ubiquitin antibodies. A strong ubiquitin-positive signal in the higher molecular range occurred upon increased expression of HOIP and HOIL-1, which was not present in cells overexpressing OTULIN, confirming a linear ubiquitin chain topology (Fig. 2D). Compared to wildtype (WT) HOIP, only a weak ubiquitin-positive signal was seen in cells overexpressing catalytically inactive HOIP C885A (Fig. 2D), which can be explained by the ability of co-expressed HOIL-1L to increase endogenous LUBAC activity. An *in vitro* ubiquitination assay using catalytically active recombinant HOIP and PINK1 immunoprecipitated from cells also supported the notion that HOIP can ubiquitinate PINK1 (Fig. 2E). We then tested for ubiquitination of endogenous PINK1 after TNF treatment. PINK1 was immunoprecipitated from untreated or TNF-treated cells and analyzed by immunoblotting using PINK1-and M1-ubiquitin-specific antibodies. Both blots showed increased signals in the high molecular weight range in TNF-treated cells which disappeared when the cell lysates were treated with recombinant OTULIN (Fig. 2F). In conclusion, TNF signaling induces recruitment of LUBAC to mitochondria, where PINK1 is stabilized.

### PINK1 counteracts OTULIN activity at mitochondria by phosphorylating M1-linked ubiquitin chains

What could be the functional consequence of PINK1 stabilization at the mitochondrial outer membrane? PINK1-mediated phosphorylation of ubiquitin alters the structure and surface properties of ubiquitin and thereby can affect the assembly and disassembly of polyubiquitin chains (Wauer et al., 2015). It has been shown previously *in vitro* that M1-linked tetra-ubiquitin is less efficiently hydrolyzed by OTULIN in the presence of PINK1 (Wauer et al., 2015). We confirmed these results (Suppl. Fig. 3A, B) and then tested whether PINK1 impairs OTULIN activity also in cells. Proteins modified by linear polyubiquitin chains were affinity-purified using the recombinant UBAN domain of NEMO (Hadian et al., 2011; Komander et al., 2009; Muller-Rischart et al., 2013; Rahighi et al., 2009), which shows a 100-fold higher affinity for M1-linked than for K63-linked ubiquitin (Lo et al., 2009; Rahighi et al., 2009), and then analyzed by immunoblotting using M1-ubiquitin-specific antibodies. The M1-ubiquitin-positive signals in extracts prepared from cells expressing HOIP and HOIL-1L were abolished by co-expression of WT OTULIN (Fig. 3A, lanes 1 and 4) but not by the inactive OTULIN variant W96A (Fig. 3A, lane 6). Co-expression of PINK1 increased M1-ubiquitination induced by HOIP and HOIL-1L (Fig. 3A, lanes 1 and 2) and partially restored M1-ubiquitination in cells co-expressing WT OTULIN (Fig. 3A, lanes 3 and 4). These results indicated that also in cells PINK1 counteracts the activity of OTULIN to hydrolyze M1-linked polyubiquitin chains.

**Figure 3.**
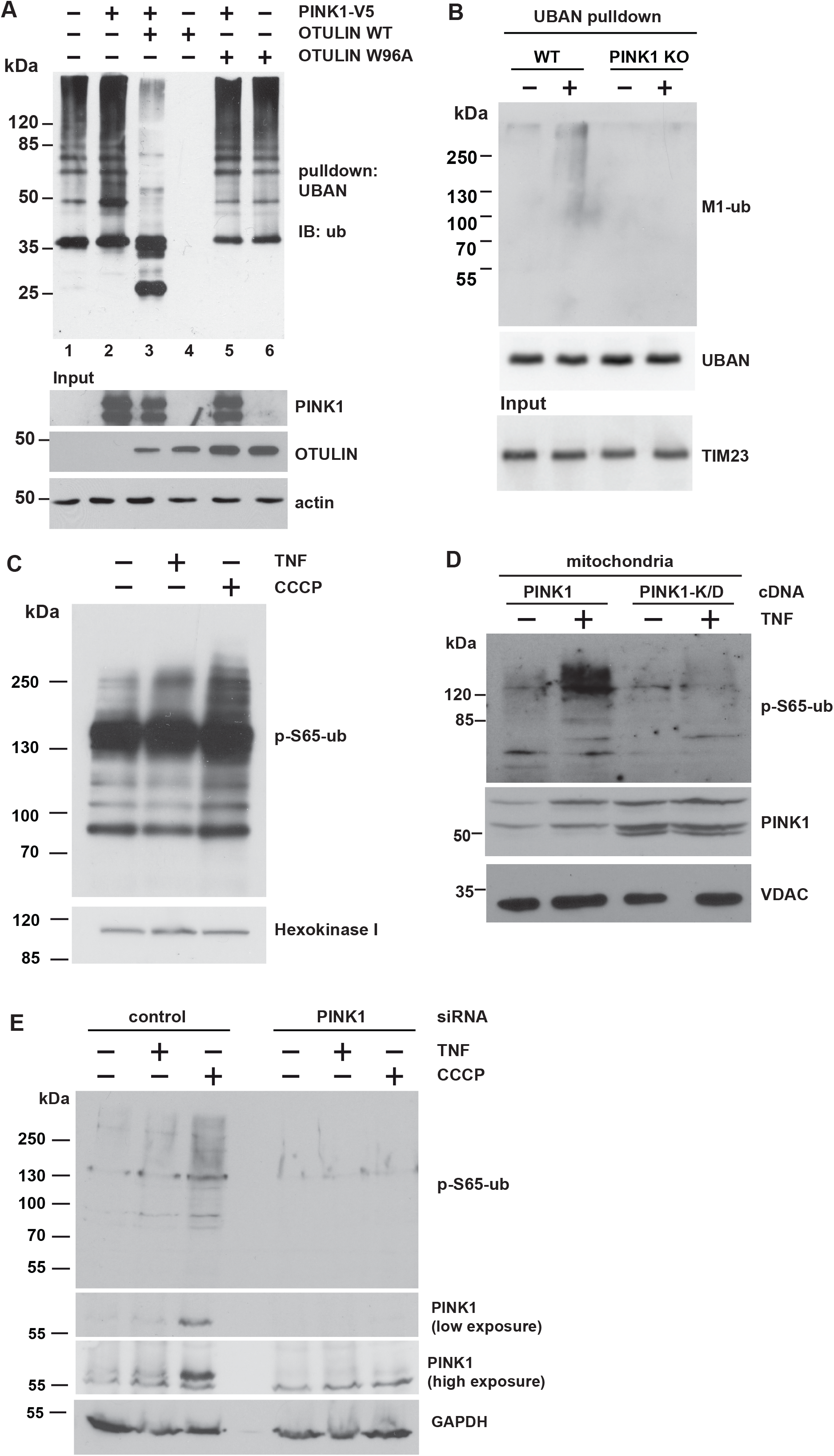
PINK1 stabilizes M1-linked ubiquitin by phosphorylation and counteracts OTULIN activity. **A. PINK1 antagonizes OTULIN activity in cells**. HEK293T cells were co-transfected with HOIP, HOIL, PINK1 and WT OTULIN or an inactive OTULIN mutant (OTULIN W96A), as indicated. The cells were lysed 24 h later under denaturing conditions, and lysates were subjected to affinity purification using the Strep-tagged UBAN domain of NEMO to enrich proteins modified with M1-linked ubiquitin. Proteins affinity-purified by Strep-Tactin beads were analyzed by immunoblotting using ubiquitin antibodies. **B. TNF-induced M1 ubiquitination is reduced in PINK1-deficient cells**. WT and PINK1-KO MEFs were treated with TNF (25 ng/ml, 15 min) and then harvested. Purified mitochondrial fractions were subjected to affinity purification using the Strep-tagged UBAN domain, as described in A. For immunoblotting, M1-ubiquitin-specifc antibodies were used. **C. TNF increases p-S65-ubiquitin**. HEK293T cells were treated with TNF (25 ng/ml, 15 min) or CCCP (10 µM, 90 min) and directly lysed in Laemmli sample buffer. Lysates were subjected to immunoblotting using p-S65-ubiquitin antibodies. The input was immunoblotted for Hexokinase 1. **D. Catalytically active PINK1 increases p-S65-ubiquitin at mitochondria**. HEK293T cells were transfected with WT PINK1 or the kinase-dead (K/D) PINK1 mutant. Two days after transfection, the cells were treated with TNF (25 ng/ml, 15 min) and lysed. Purified mitochondrial fractions were analyzed by immunoblotting using antibodies against p-S65-ubiquitin and PINK1. The input was immunoblotted for VDAC. **E. The TNF-induced increase in p-S65-ubiquitin at mitochondria is abolished in PINK1-deficient cells**. HEK293T cells were transfected with control or PINK1-specific siRNA and treated with TNF (25 ng/ml, 15 min) or CCCP (10 µM, 90 min) two days after transfection. Purified mitochondrial fractions were analyzed as described in D.

Prompted by these findings, we wondered whether PINK1 increases TNF-induced M1-ubiquitination at mitochondria by phosphorylating ubiquitin. TNF-induced M1-linked ubiquitination was indeed reduced in MEFs from PINK1 KO mice (Fig. 3B). We also observed an increased abundance of p-S65-ubiquitin within 15 min of TNF treatment, albeit less pronounced than in the positive control (CCCP, 90 min) (Fig. 3C), which is consistent with the lower amount of PINK1 stabilized by TNF (Fig. 2A). Moreover, ectopic expression of WT PINK1 but not kinase-dead (K/D) PINK1 increased p-S65-ubiquitin in the mitochondrial fraction after TNF treatment (Fig. 3D), whereas in cells silenced for PINK1 expression TNF did not increase p-S65-ubiquitin (Fig. 3E).

### Mitochondrial M1 ubiquitination protects from apoptosis

Our data so far revealed that TNF increases M1-linked ubiquitination at mitochondria and stabilizes PINK1 without inducing mitophagy. Based on the fact that both TNF and PINK1 have anti-apoptotic functions, we were wondering whether M1-ubiquitination at mitochondria protects from cell death. TNF induces the increased expression of anti-apoptotic proteins through formation of complex I at the TNFR1 and activation of the transcription factor NF-ĸB (Brenner et al., 2015; Hayden and Ghosh, 2014; Kupka et al., 2016b). We hypothesized that the fast TNF-induced increase in mitochondrial ubiquitination protects mitochondria from apoptosis before transcriptional reprogramming by NF-ĸB takes effect. To address this possibility experimentally, we exposed cells to a short pro-apoptotic stimulus with or without TNF pretreatment for 15 min. Apoptotic cell death was assessed by mitochondrial Bax recruitment, cytochrome c release, caspase-3 activation, and an ultrastructural analysis of cristae morphology. Bax was efficiently recruited to mitochondria within 1 h of staurosporine (STS) treatment, as detected by immunoblotting of purified mitochondria (Fig. 4A). When STS was applied 15 min after TNF pretreatment, the amount of mitochondrial Bax was reduced (Fig. 4A). Likewise, STS-induced cytochrome c release into the cytoplasm was decreased by the TNF pretreatment (Fig. 4B). Electron microscopy revealed that TNF prevents the STS-induced reduction in cristae abundance and cristae length per mitochondrial area (Fig. 4C, D). Moreover, TNF significantly reduced the number of STS-treated cells with activated caspase-3 (Fig. 4E). This effect was not compromised in the presence of the NF-ĸB super-repressor IĸBα (serines 32 and 36 replaced by alanines), which blocks nuclear translocation of the NF-κB subunit p65 (Sun et al., 1996), indicating that the fast protective effect of TNF was not mediated by NF-ĸB activation (Fig. 4E). Notably, both HOIP and PINK1 were required for the protective effect of TNF pretreatment, since in cells silenced for HOIP or PINK1 expression, TNF was not able to reduce STS-induced cytochrome c release (Fig. 4F, G). We concluded that the TNF-induced assembly of M1-linked ubiquitin by HOIP and their phosphorylation by PINK1 interferes with the insertion of Bax into the outer mitochondrial membrane, thereby preventing apoptotic cell death. To check for mitochondrial fitness, we also analyzed cellular bioenergetics in response to TNF treatment and observed a transient increase in maximal respiration, spare respiratory capacity and ATP production (Suppl. Fig. 4A-C), suggesting that TNF enhances energy metabolism.

**Figure 4.**
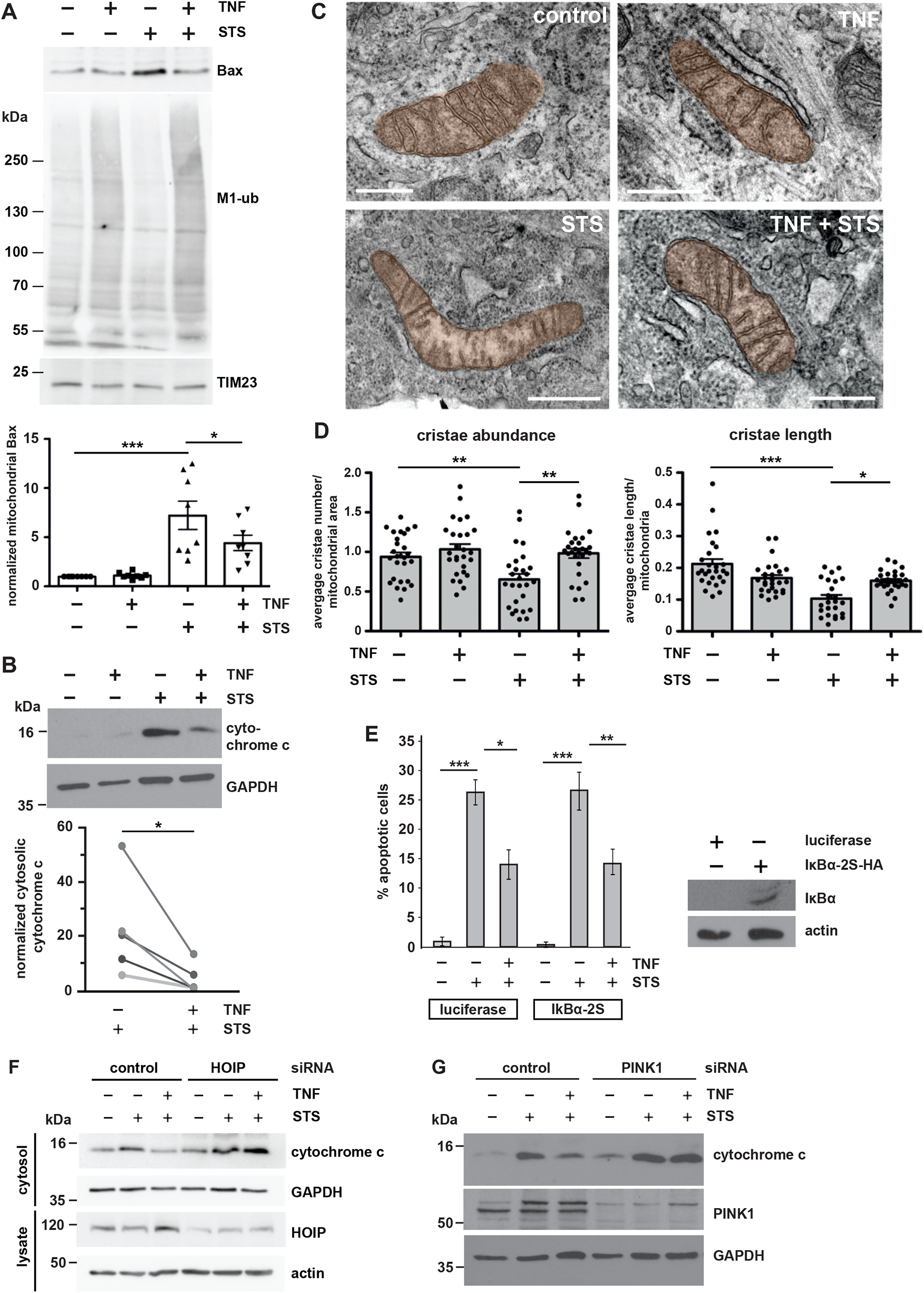
M1 ubiquitination at mitochondria protects from apoptosis. **A. Mitochondrial Bax recruitment is reduced by TNF**. HeLa cells were treated with STS (1 μM, 1 h) with or without a 15 min pretreatment with TNF (25 ng/ml) and then harvested. Purified mitochondrial fractions were analyzed by immunoblotting using antibodies against Bax and M1-ubiquitin. The input was immunoblotted for TIM23 (upper panel). Bax-specific signal intensities were quantified and normalized to TIM23-specific signals (lower panel). Data represent the mean with standard error of 8 independent experiments. *p < 0.05, ***p < 0.001. Statistics: Kruskal-Wallis test. **B. Cytochrome C release is reduced by TNF**. HeLa cells were treated as described in A. The cytosolic fractions were analyzed by immunoblotting using cytochrome c antibodies. GAPDH was used as a reference. Quantification of 5 biological replicates is shown in the lower panel. Signal intensities were quantified and normalized to that of GAPDH. *p < 0.05. **C, D. TNF treatment prevents damage of mitochondrial cristae under pro-apoptotic conditions**. C. SH-SY5Y cells were treated as described in A, fixed, embedded and the mitochondrial ultrastructure was imaged by electron microscopy. Scale bar: 400 nm. D. Cristae abundance (average cristae number per mitochondrial area) and cristae length (average cristae length per mitochondrium) were analyzed by Imaris 9.8. At least 27 mitochondria were assessed per condition. *p < 0.05, **p < 0.01, ***p < 0.01. **E. The fast anti-apoptotic effect of TNF is not affected by the NF-κB inhibitor IκBα**. SH-SY5Y cells were transiently transfected with the NF-κB super-repressor IκBα-2S or luciferase as a control. One day after transection, the cells were treated with STS (5 µM, 2 h) with or without a 15 min pretreatment with TNF (25 ng/ml). Cells were fixed and stained by active caspase-3 antibodies. Signal intensities were quantified by immunocytochemistry and fluorescence microscopy. Data represent the standard deviation of three independent experiments; at least 300 cells were counted per experiment. *p < 0.05, **p < 0.005. Expression of IκBα-2S was tested by immunoblotting using antibodies against the HA tag. **F. The protective effect of TNF is dependent on HOIP expression**. HeLa cells were transiently transfected with control or HOIP siRNA. Two days after transfection, cells were treated with STS (1 μM, 1 h) with or without a 15 min TNF pretreatment (25 ng/ml) and then harvested. The cytosolic fractions were analyzed by immunoblotting using cytochrome c antibodies. HOIP silencing efficiency was analyzed in whole cell lysates using antibodies against HOIP. GAPDH and actin were immunoblotted as input controls. **G. The protective effect of TNF is dependent on PINK1 expression**. HeLa cells were transiently transfected with control or PINK1 siRNA and treated as described in F. The cytosolic fractions were analyzed by immunoblotting using cytochrome c and PINK1 antibodies. GAPDH was immunoblotted as input control.

### A signaling platform for TNF-induced NF-ĸB activation is assembled at mitochondria

Linear ubiquitin chains specifically interact with proteins harboring an UBAN domain, which is found for example in NEMO and Optineurin. NEMO is the regulatory subunit of the IκB kinase (IKK) complex that not only binds to M1-linked ubiquitin chains but is also a key substrate of HOIP for linear ubiquitination. NEMO ubiquitination and subsequent oligomerization activates the associated kinases IKKα and IKKβ, required for NF-κB activation. Though the fast TNF-mediated protection of mitochondria from apoptosis was NF-κB-independent, we observed an increased abundance of NEMO, p65, and phosphorylated p65 (p-65) at mitochondria after 15 min TNF treatment (Fig. 5A). We therefore speculated that mitochondrial M1-ubiquitination contributes to TNF signal amplification and facilitates transmission of the signal to the nucleus. We first validated our results in primary human macrophages, a cell type highly relevant for TNF signaling. Interestingly, a fraction of NEMO and p65 resided at mitochondria already in non-stimulated macrophages. Phosphorylation of p65 increased after TNF treatment (Fig. 5B), suggesting the formation of a local NF-κB signaling platform at mitochondria. Next, we tested whether PINK1 has a role in mitochondrial NF-κB pathway activation. TNF-induced nuclear translocation of p65 was significantly decreased in cells silenced for PINK1 expression. This phenotype was rescued by WT PINK1 but not by the catalytically inactive PINK1-K/D mutant (Fig. 5C). Since NEMO is a highly relevant substrate of HOIP in NF-κB pathway activation, we explored the possibility that PINK1 phosphorylates ubiquitinated NEMO. In cells expressing WT PINK1, the p-S65-ubiquitin signal was strongly enhanced at NEMO affinity-purified from cell lysates, whereas no signal was seen when catalytically inactive PINK1-K/D was expressed (Fig. 5D). Supporting the physiological relevance of endogenous PINK1, TNF-induced M1-ubiquitination of NEMO was reduced in PINK1-deficient cells (Fig. 5E). Taken together, catalytically active PINK1 promotes NF-κB activation, most likely by stabilizing M1-linked ubiquitin chains through phosphorylation.

**Figure 5.**
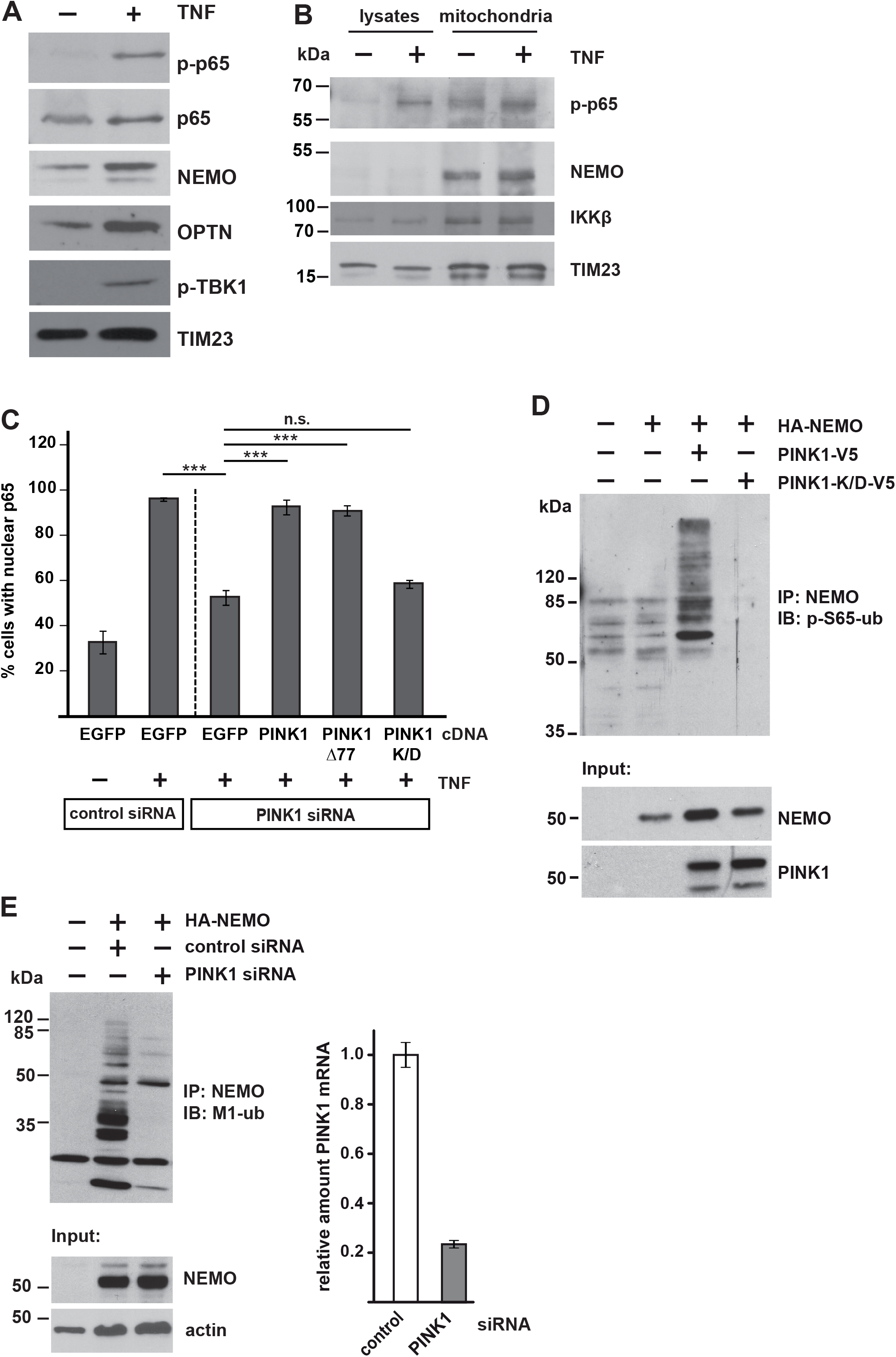
Mitochondria serve as a signaling platform for TNF-induced NF-ĸB activation. **A, B. NF-κB signaling components are recruited to mitochondria upon TNF treatment.** A. HEK293T cells were treated with TNF (25 ng/ml, 15 min), harvested and purified mitochondrial fractions were analyzed by immunoblotting using the antibodies indicated. B. Primary human macrophages were treated and analyzed as described in A. Whole cell lysates (5%) were analyzed in parallel to mitochondrial fractions (30%) using the indicated antibodies. **C. Nuclear translocation of p65 is impaired in cells silenced for PINK1 expression**. SH-SY5Y cells were transfected with control or PINK1-specific siRNAs. For rescue experiments, cells were co-transfected with WT PINK1, PINK1Δ77, or kinase-dead PINK1-K/D. Two days after transfection the cells were treated with TNF (25 ng/ml, 25 min), fixed and stained with p65 antibodies. The fraction of cells showing nuclear translocation of p65 was determined for each condition. Data represent the mean ± SEM of 3 independent experiments. The experiment was performed in triplicates and more than 300 cells were quantified per experiment and condition. ***p < 0.001. **D. PINK1 phosphorylates ubiquitinated NEMO**. HEK293T cells were co-transfected with V5-tagged WT PINK1 or kinase-dead PINK1-K/D and HA-tagged NEMO. One day after transfection the cells were lysed and subjected to immunoprecipitation using antibodies against HA. Precipitated proteins were then detected by immunoblotting using p-S65-ubiquitin antibodies. The input was immunoblotted for NEMO and PINK1. **E. Linear ubiquitination of NEMO is reduced in PINK1-deficient cells**. HEK293T cells were co-transfected with HA-NEMO and control or PINK1-specific siRNAs. Two days after transfection the cells were lysed and subjected to immunoprecipitation using antibodies against HA. Precipitated proteins were then detected by immunoblotting using M1-ubiquitin antibodies. The input was immunoblotted for NEMO and PINK1. PINK1 silencing efficiency was determined by real-time RT-PCR.

### TNF induces shuttling of p65 to the nucleus and increases mitochondria-nucleus contact sites

What could be the advantage of assembling a signaling platform at mitochondria? First, the mitochondrial network provides a large surface area for signal amplification. Second and possibly even more relevant, mitochondria are highly dynamic and mobile organelles that engage in physical and functional interactions with other organelles and the plasma membrane (Giacomello et al., 2020; Harper et al., 2020 ; Prinz et al., 2020; Scorrano et al., 2019). We therefore wondered whether mitochondria facilitate the transport of activated NF-κB components to the nucleus. First, we followed up the motility of mitochondria in primary human macrophages upon TNF stimulation by life cell imaging and observed a decrease in the distance between peripheral mitochondria and the nucleus within 15 min of TNF treatment (Fig. 6A and Movie 1). Encouraged by this observation, we tested whether mitochondrial motility affects the nuclear translocation of p65 upon TNF stimulation. We silenced the expression of the mitochondrial outer membrane Rho GTPases Miro1 and Miro2, which are components of the adaptor complex that anchors mitochondria to motor proteins (Misgeld and Schwarz, 2017; Schwarz, 2013). A ratiometric analysis revealed a significant decrease in the fraction of nuclear p65 upon TNF stimulation in Miro1/2-deficient cells (Fig. 6B). We then reasoned that mitochondrial transport of p65 to the nucleus should increase the proximity of mitochondria to the nucleus and possibly favor the formation of mitochondria-nucleus contact sites. To this end, we have generated a new genetically encoded split-GFP contact site sensor (SPLICS) (Cali and Brini, 2021; Cieri et al., 2018; Vallese et al., 2020), capable to reconstitute fluorescence only when the nucleus and mitochondria are in close proximity (Fig. 6C). Correct targeting and self-complementation ability of the GFP_1-10_ and the β11 split-GFP fragments to the outer nuclear membrane (ONM) were tested (Suppl. Fig. 5A-C), and the ONM-β11 was further selected to generate a SPLICS reporter along with the outer mitochondrial membrane (OMM) targeted GFP_1-10_, extensively used in all mitochondrial sensors generated so far. Therefore, a first generation (SPLICS^NU-MT^) (Cieri et al., 2018) and a second generation (SPLICS-P2A^NU-MT^) of reporters improved to express equimolar amounts of the organelle-targeted GFP fragments in a bicistronic vector (Kim et al., 2011; Vallese et al., 2020) was generated (Fig. 6D). Expression of the SPLICS^NU-MT^ in HeLa cells resulted in the emission of a fluorescent punctate signal within the nuclear/perinuclear region that could be easily quantified with the Fiji (https://imagej.net/software/fiji/) software, and custom macros written for interorganelle contacts analysis (Cali and Brini, 2021), with an additional ROI traced around the nucleus of SPLICS^NU-MT^-positive cells to specifically focus only contacts that involve the nuclear envelope (Suppl. Fig. 5D). We have tested the ability to detect changes in the nucleus-mitochondria contact sites by triggering the mitochondrial retrograde response (MRR) in HeLa cells expressing the SPLICS^NU-MT^ reporter (Amuthan et al., 2001; Desai et al., 2020; Eisenberg-Bord and Schuldiner, 2017; Walker and Moraes, 2022). Accordingly, MRR activation induced an increase in the nucleus-mitochondria contact sites (Suppl. Fig. 5E). With this new tool we assessed whether mitochondrial transport of p65 to the nucleus favors the proximity of mitochondria to the nucleus by TNF treatment of SPLICS^NU-MT^ expressing cells. Indeed, the number of mitochondria-nucleus contact sites strongly increased upon TNF stimulation (Fig. 6E-G). Remarkably, HOIP silencing by siRNA completely abolished the TNF-induced increase of the mitochondria-nucleus contacts (Fig. 6H-J). Thus, all these approaches supported the notion that mitochondria are implicated in shuttling activated NF-κB to the nucleus upon TNF stimulation and that this process is dependent on linear ubiquitination.

**Figure 6.**
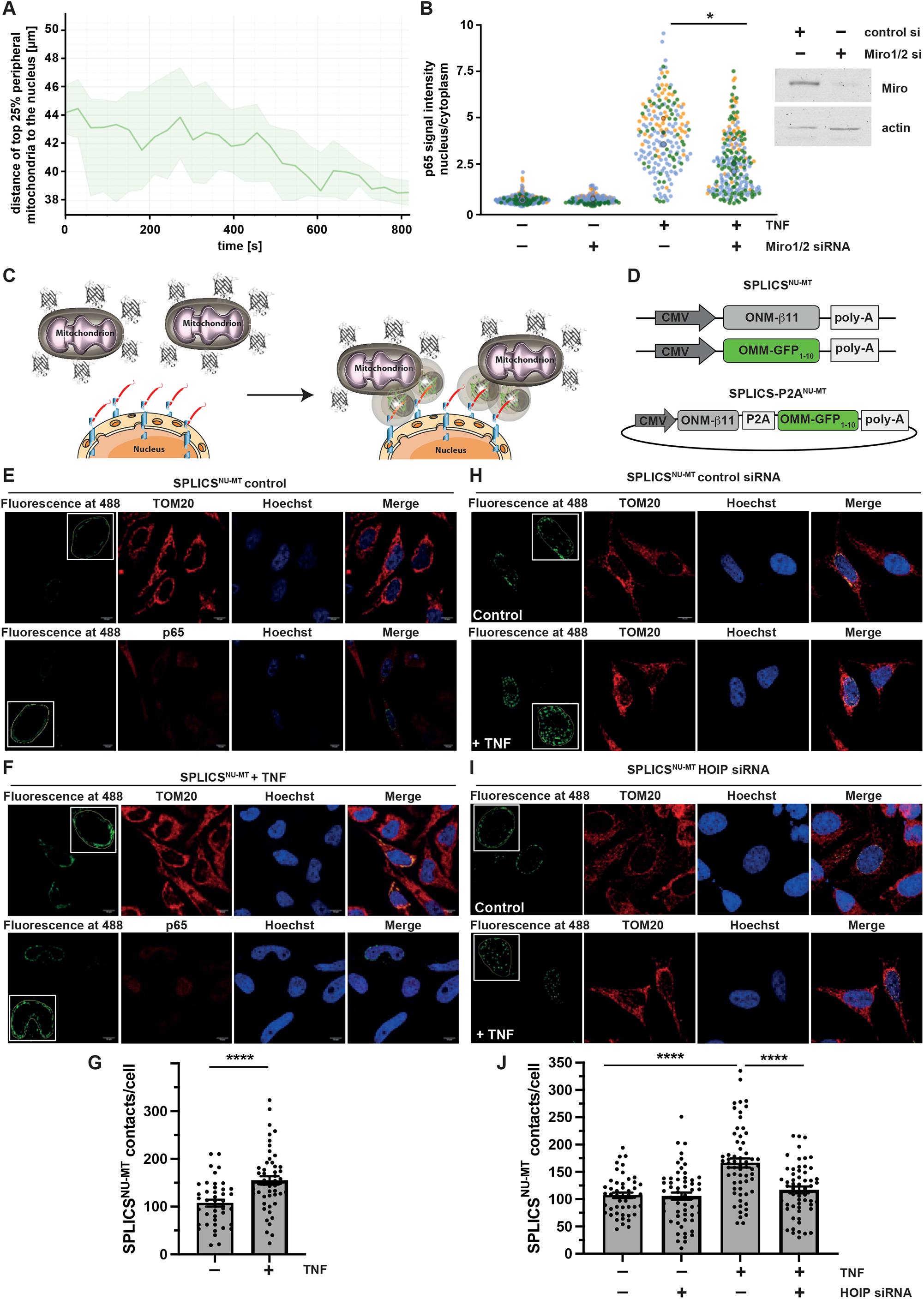
Mitochondria facilitate the transfer of p65 to the nucleus. **A. TNF induces the movement of peripheral mitochondria towards the nucleus**. Primary macrophages were stained by Mitotracker™ Green to visualize mitochondria and by Hoechst 33342 to visualize the nucleus and monitored every 30 s for 15 min after treatment with TNF (25 ng/ml). The motility of the mitochondria was analyzed using Imaris 9.8 spot function. The graphical representation of the top 25% peripheral mitochondrial indicates a decrease of the average mitochondrial distance to the nucleus after TNF treatment. **B. Nuclear p65 translocation upon TNF treatment is reduced cells silenced for Miro1/2 expression**. HeLa cells were transfected with control or Miro1 and Miro2 siRNAs. Two days later, cells were treated with TNF (25 ng/ml, 15 min), fixed and stained using antibodies against p65 and tubulin and DAPI. Images were segmented in a cytoplasmic and nuclear compartment and the fluorescence signal intensity ratio of p65 was measured using CellProfiler 4.2.1. Quantification is based on 3 biological replicates. Per condition at least 79 cells have been analyzed. ***p < 0.001. **C. Cartoon of the newly developed SPLICS for the detection nucleus-mitochondria contacts**. Fluorescence reconstitution between the ONM-targeted β11 and the OMM-targeted GFP_1-10_ fragment occurs at the contact site. **D. Schematic depiction of the plasmids encoding the 1**^**st**^ **generation (SPLICS**^**NU-MT**^**) and the 2**^**nd**^ **generation (SPLICS-P2A**^**NU-MT**^**) reporters**. **E, F, G. TNF increases mitochondria-nucleus contact sites**. Representative immunofluorescence images (E, F) and quantification (G) of mitochondria-nucleus contacts in HeLa cells transfected with the SPLICS^NU-MT^ and treated with 25 ng/ml TNF or PBS for 15 minutes. TOM20 antibodies were used to stain mitochondria and p65 antibodies to assess p65 nuclear translocation. Nuclei were stained with Hoechst33342 (ThermoFisher, 1 μg/ml). **H, I, J. TNF increases mitochondria-nucleus contact sites in a HOIP-dependent manner**. Representative immunofluorescence images (H, I) and quantification (J) of mitochondria-nucleus contacts in control or HOIP-silenced (HOIP siRNA) HeLa cells transfected with the SPLICS^NU-MT^ probe treated with 25 ng/ml TNF or PBS for 15 minutes. Mitochondria and nuclei were stained as described in E and F.

## DISCUSSION

The formation of mitochondrial signaling platforms has been implicated in various signaling paradigms. However, many aspects of why cells employ mitochondria as signaling organelles are still unknown. One obvious advantage is that the integration of mitochondria in signaling pathways facilitates context-dependent adaptive responses to bioenergetic or biosynthetic demands. Another plausible explanation is that several signaling pathways affect cell fate decisions, which are regulated by mitochondria. Our study revealed two other advantages of mitochondrial signaling platforms. Mitochondria can contribute to signal amplification by virtue of their large surface and the presence of signaling components that stabilize “signals” by forming protein complexes (signalosomes) and/or by posttranslational modifications of signaling components. In TNF signaling, the mitochondrial kinase PINK1 performs such a function. It increases the stability of M1-linked ubiquitin chains assembled by HOIP by antagonizing the M1-ubiquitin-specific deubiquitinase OTULIN, thus enhancing downstream signaling. Moreover, mitochondria are highly dynamic and mobile organelles, allowing them to communicate and interact with various cellular components and organelles. Here we present evidence that mitochondria can act as vehicles to shuttle activated transcription factors to the nucleus. Efficient nuclear translocation of the NF-κB subunit p65 upon TNF stimulation was impaired in cells silenced for Miro1 and Miro2 expression. Even more striking, we found that TNF increases mitochondria-nucleus contact sites, which presumably facilitate the uptake of p65 into the nuclear compartment. In support of this notion, the expression level of a mitochondria-nucleus tether component identified in breast cancer cells, the mitochondrial translocator protein TSPO, correlated with the abundance of nuclear NF-κB (Desai et al., 2020). TNF-induced formation of mitochondria-nucleus contact sites was dependent on HOIP, confirming a role of M1-linked ubiquitination. In a recent study performed in *Saccharomyces cerevisiae*, TOM70 has been implicated in the formation of tethers between mitochondria and the nucleus (Eisenberg-Bord et al., 2021). TOM70 has also been identified as a target for ubiquitination within the TNF signaling network (Wagner et al., 2016) and was detected as an OTULIN-interacting protein in our mass spectrometry-based screen. In addition, TOM70 is required for the activation of IRF3 in MAVS signaling (Liu et al., 2010; Thorne et al., 2022). Hence, it will be interesting to explore a possible role of TOM70 in the NF-κB pathway.

TNF signaling usually mounts an anti-apoptotic response via the TNFR1/complex I/NF-κB axis, implicating transcriptional regulation of anti-and pro-apoptotic gene expression (Varfolomeev and Vucic, 2018). We previously identified the mitochondrial GTPase OPA1 as an NF-κB target gene, which contributes to anti-apoptotic reprogramming by maintaining cristae integrity (Muller-Rischart et al., 2013). Here we identified a fast, transcription independent anti-apoptotic TNF response that protects mitochondria by remodeling the outer mitochondrial membrane. This remodeling involves the assembly of M1- and K63-linked ubiquitin chains and the recruitment of ubiquitin-binding proteins and signaling components to the outer mitochondrial membrane, which obviously impedes Bax insertion into the outer membrane under pro-apoptotic conditions. Conceptually, the fast TNF response ensures mitochondrial integrity and fitness to fulfil their role in signal amplification and transport of activated NF-ĸB to the nucleus. Along this line, we observed that TNF increases maximal respiration and spare respiratory capacity within 15 min.

Finally, our study unveiled a mitophagy-independent function of PINK1 in mitochondrial signaling. In contrast to mitophagy-inducing conditions, PINK1 is stabilized in TNF signaling without dissipation of the mitochondrial membrane potential. Accordingly, proteolytically processed, mature PINK1 is stabilized in addition to full-length PINK1 destined for import through the TOM/TIM23 complexes. Albeit the accumulation of the mature PINK1 species after 15 min TNF treatment is less striking than that of unprocessed PINK1 after 90 min CCCP treatment, several lines of evidence point towards a physiological role of PINK1 in NF-ĸB signaling. First, PINK1 phosphorylates M1-linked ubiquitin chains and thereby decreases the efficiency of OTULIN to hydrolyze linear ubiquitin chains. This explains why TNF-induced M1-ubiquitination is decreased in PINK1-deficient cells. Second, the fast anti-apoptotic TNF response depends on both HOIP and PINK1 expression. PINK1 is stabilized by complex formation with HOIP and HOIP-dependent linear ubiquitination and in turn augments M1-ubiquitination by antagonizing their disassembly. This scenario is reminiscent of the feed-forward activation loop in PINK1/Parkin-dependent mitophagy and completes the picture of a mitophagy-independent function of PINK1 and Parkin under physiological conditions that integrates their roles in innate immune signaling and stress protection.

## ACKNOWLEDGEMENT

We thank Jean-François Trempe for providing the tcPINK1 plasmid, Ben Stieglitz for the HOIP-RBR-LDD and UBE2L3 plasmids, Daniel Krappmann for the OTULIN plasmids, and Sadasivam Jeganathan for protein expression and purification. KFW is supported by the German Research Foundation (WI/2111-4, WI/2111-6, WI/2111/8, FOR 2848, and Germany’s Excellence Strategy - EXC 2033 - 390677874 – RESOLV) and the Michael J-Fox Foundation for Parkinson’s Research (Grant ID 16293). SR-SIM microscopy was funded by the German Research Foundation and the State Government of North Rhine-Westphalia (INST 213/840-1 FUGG). TC is supported by grants from the Ministry of University and Research (Bando SIR 2014 no. RBSI14C65Z and PRIN2017) and from the Università degli Studi di Padova (Progetto Giovani 2012 no. GRIC128SP0 to T.C., Progetto di Ateneo 2016 no. CALI_-SID16_01, STARS Consolidator Grant 2019). M.B is supported by Local Funds from the University of Padova. CM is supported by the German Research Foundation (Project-ID 390339347, Emmy Noether Programme and Project-ID 259130777, CRC1177). WS is supported by the National Institutes of Health (NS085070, NS110085, and NS110435), the Department of Defense Congressionally Directed Medical Research Programs (W81XWH-17-1-0248), the Michael J. Fox Foundation for Parkinson’s Research, Mayo Clinic Foundation and the Center for Biomedical Discovery (CBD).

---

**Suppl. Figure 1. Identification of OTULIN-interacting proteins by mass spectrometry S1A. OTULIN-interacting proteins**. Proteins identified by co-immunoprecipitation coupled to mass spectrometry analysis using strict filter criteria. Only proteins that were not identified in the control and that were reliably identified in both OTULIN co-immunoprecipitations by at least two unique peptides are listed according to their abundance (iBAQ-value). Mitochondria-associated proteins are marked with asterisk.

**S1B. Gene Ontology cellular compartment (GOCC) characterization of OTULIN-interacting proteins**. The GOCC classifications for each of the identified proteins were performed using DAVID.

**Suppl. Figure 2. PINK1 interacts with the N-terminal part of HOIP including the UBA domain**

**S2A, C. Schematic representation of the HOIP constructs used to map the interaction between HOIP and PINK1**. All constructs are equipped with an N-terminal HA-tag. PUB: peptide N-glycosidase/ubiquitin-associated domain; ZF: zinc finger domain; NZF: nuclear protein localization 4-type zinc finger domain; UBA: ubiquitin-associated domain; RING: really interesting new gene; IBR: in-between RING domain; LDD: linear ubiquitin chain determining domain.

**S2B. PINK1 co-immunoprecipitates with the N-terminal part of HOIP1**. HEK293T cells were transfected with V5-tagged PINK1 and the indicated HA-tagged HOIP constructs. One day after transfection cells were lysed under native conditions and subjected to immunoprecipitation using HA antibodies. Immunopurified proteins were analyzed by immunoblotting using V5 antibodies to detect PINK1.

**S2D. The UBA domain enhances the interaction between HOIP and PINK1**. HEK293T cells were analyzed as described in B.

**S2E. PINK1 co-elutes with HOIP in cellular lysates**. HEK293T cells expressing V5-tagged PINK1 were lysed, and soluble proteins were separated by size-exclusion chromatography. Fractions were collected and analyzed by immunoblotting using antibodies against HOIP and V5.

**Suppl. Figure 3. PINK1 impairs OTULIN activity *in vitro***

**S3A. The efficiency of OTULIN to hydrolyze M1-linked ubiquitin is decreased in the presence of PINK1**. Recombinant M1-linked tetra-ubiquitin was incubated with or without recombinant *Tribolium castaneum* tcPINK1 for 48 h. Then recombinant OTULIN was added for the indicated time. The reaction was stopped by adding Laemmli sample buffer and the samples were analyzed by immunoblotting using ubiquitin antibodies.

**S3B. PINK1 phosphorylates M1-linked tetra-ubiquitin *in vitro***. M1-linked tetra-ubiquitin was incubated with or without recombinant tcPINK1 in kinase buffer for 2 days. The samples were analyzed by Phos-tag™ SDS-PAGE and immunoblotting using ubiquitin antibodies.

**Suppl. Figure 4. TNF influences the bioenergetic profile**

**S4A, B. TNF increases maximal respiration (A) and spare respiratory capacity (B)**. HeLa cells were treated with TNF for the indicated time and analyzed by the Agilent Seahorse XF Cell Mito Stress Test. The spare respiratory capacity is the difference between the maximal and the basal respiration.

**S4C. TNF TNF increases total ATP production**. HeLa cells were treated with TNF for indicated time and analyzed by the Agilent Seahorse ATP Rate Assay.

**Suppl. Figure 5. Targeting, self-complementation and ability to respond to the MRR activation of the SPLICS**^**NU-MT**^ **reporters**.

**S5A**. Assessment of the nuclear envelope localization of the ONM-targeted GFP_1-10_ fragment at different concentrations of plasmid transfected in HeLa cells. The GFP_1-10_ fragment was stained with a mouse anti-GFP primary antibody (Santa Cruz, 1:100) followed by an anti-mouse AlexaFluor 488-conjugated secondary antibody (ThermoFisher, 1:100).

**S5B**. Assessment of the self-complementation capability of the ONM targeted GFP_1-10_ fragment with a cytosolic mKate-tagged β11.

**S5C**. Nuclear envelope localization and self-complementation of the ONM-targeted β11 fragment co-transfected with an untargeted, cytosolic GFP1-10 fragment. Nuclei were stained with Hoechst33342 (ThermoFisher, 1 μg/ml).

**S5D**. Schematic representation of the contact quantification process using ImageJ plugin VolumeJ and custom-made macros specifically developed for the analysis: 1. a ROI is traced around the nucleus of the cell; 2. the first macro is applied, which clears all the signal outside the ROI and convolves the signal inside; 3. VolumeJ is used to generate a rendering of the convolved signal; 4. the second custom macro is applied on the first and the last frame of the rendering; 5. the contacts per each frame are quantified and the overall number of contacts per cell is calculated by averaging the two results.

**S5E**. Representative images and contact quantification of HeLa cells after induction of the mitochondrial retrograde response (MRR) upon treatment with rotenone (5 µM for 3 h), CCCP (10 µM for 3 h) or staurosporine (1 µM for 2 h). Mitochondria were stained with MitoTracker Red CMXRos (ThermoFisher, 100 nM) in HBSS for 30 minutes. Nuclei were stained as described in C.

**Movie 1. TNF induces the movement of peripheral mitochondria towards the nucleus**. SH-SY5Y cells were stained with NucSpot Live 488 and MitoView 650 for 30 min. After TNF treatment (25 ng/ml) the cells were imaged every 2 min for 30 min. Image analysis was performed by Imaris 9.8 using the Surface (nucleus) and Spot (mitochondria) algorithm. Mitochondrial distance with respect to the nuclear surface was color-coded (purple/blue = proximal; red/yellow =distal).

## METHODS

### Cell culture

#### HEK293T, HeLa cells, Mouse embryonic fibroblasts (MEFs)

Cells were cultured in Dulbecco’s modified Eagle’s medium (DMEM) supplemented with 10% (v/v) fetal bovine serum (FBS) and 100 IU/ml penicillin and 100 μg/ml streptomycin sulfate.

#### SH-SY5Y cells

Cells were cultured in Dulbecco’s modified Eagle’s medium F-12 (DMEM/F12) supplemented with 15% (v/v) fetal bovine serum (FBS), 100 IU/ml penicillin, 100 μg/ml streptomycin sulfate and 1% non-essential amino acids.

#### HAP1 WT and HAP1 HOIP (RNF31) KO

Cells were cultured in Iscove’s Modified Dulbecco’s Medium (IMDM) supplemented with 10% (v/v) FBS, 100 IU/ml penicillin, 100 μg/ml streptomycin sulfate, and 8 mM L-glutamine.

#### Human primary macrophages

Human peripheral blood mononuclear cells (PBMC) were isolated from commercially available buffy coats from anonymous donors (DRK-Blutspendedienst Baden-Württemberg - Hessen, Institut für Transfusionsmedizin und Immunhämatologie, Frankfurt, Germany) using Pancoll (PAN Biotech, Aidenbach, Germany) density centrifugation. Monocytes were separated from PBMC by adherence to plastic after 1 h incubation in serum-free RPMI1640 medium. To differentiate to macrophages, monocytes were cultured in RPMI1640 medium (ThermoFisher Scientific, Waltham, MA, USA) supplemented with 100 U/mL penicillin, 100 μg/mL streptomycin and 3% human serum (DRK-Blutspendedienst Baden-Württemberg - Hessen) for 7 days followed by culture in RPMI 1640 medium containing 10% fetal calf serum.

### Transfection and siRNA knockdown

Unless described otherwise, the transfection was performed using the following procedure: For SH-SY5Y and HeLa cells, transient transfection was performed with Lipofectamine and Plus Reagent (Invitrogen) according to the manufacturer’s instructions. For RNA interference, cells were transfected with stealth siRNA oligos (Invitrogen) using Lipofectamine RNAiMAX (Invitrogen) or Lipofectamine 2000 (Invitrogen) for co-transfection of siRNA and DNA plasmids. For HEK293T cells, constructs were transfected via PEI (polyethylenimine). DNA and PEI were mixed in a 1:2 ratio (µg: µl) in Opti-MEM and incubated for 15 min at room temperature. The DNA-PEI mixture were then added to the cells.

#### SPLICS^NU-MT^ transfection

Transient transfection was performed using the calcium phosphate method. 30,000 HeLa cells/well were seeded on a 24-well plate on coverslips in the evening. The morning after, a transfection mixture was prepared as following (quantities reported per well): 50 µl HEPES buffered saline 2x (Merck), 5 µl 2.5 M CaCl_2_, 1+1 µg SPLICSNU-MT plasmids, and up to 100 µl sterile H_2_O. The mixture was added to the wells, each containing 500 µl of growth medium. After 8 h, the mixture-containing medium was removed and replaced with fresh medium. 24 h after the medium change, the cells were treated as described below and fixed.

#### SPLICS^NU-MT^ and siRNA co-transfection

Co-transfection of SPLICS^NU-MT^ plasmids and siRNAs for HOIP silencing was performed using Lipofectamine 3000 (Thermo Fisher). 400,000 HeLa cells/well were seeded on a 6-well plate in the evening. The morning after, the medium was replaced with 750 µl/well Opti-MEM (Thermo Fisher) and a transfection mixture with the following composition per well was prepared and added to the wells: 250 µl Opti-MEM, 0.5 + 0.5 µg SPLICS^NU-MT^ plasmids, 60 µM each (180 µM in total) stealth siRNA oligos (Invitrogen), and 5 µl Lipofectamine 3000. 32 h after the addition of the transfection mixture, the cells were detached using trypsin and re-seeded on a 24-well plate on coverslips (30,000 cells/well). 16 h after the re-seeding, the cells were treated as described below and fixed.

### Immunoblotting

Proteins were size-fractionated by SDS-PAGE and transferred to nitrocellulose or polyvinylidene difluoride membranes by electroblotting. The nitrocellulose membranes were blocked with 5% nonfat dry milk or 5% BSA in TBST (TBS containing 0.1% Tween 20) for 60 min at room temperature and subsequently incubated with the primary antibody diluted in blocking buffer for 16 h at 4°C. After extensive washing with TBST, the membranes were incubated with horseradish peroxidase-conjugated secondary antibody for 60 min at room temperature. Following washing with TBST, the antigen was detected with the enhanced chemiluminescence (ECL) detection system (Promega) as specified by the manufacturer. In addition, immunoblots with fluorescently labeled secondary antibodies were imaged by an Azure Sapphire Biomolecular Imager (Azure Biosystems, USA). For quantification of Western blots, Image Studio lite (version 3.1) and Fiji was used for X-ray films and AzureSpot (version 2.0) was used for the Sapphire Biomolecular Imager (Azure Biosystems, USA).

### Mitochondrial bioenergetics

Metabolism of living cells was measured in real-time using Seahorse XFe Analyzer. The Seahorse Cell Mito Stress Test was employed to measure basal mitochondrial respiration, maximal mitochondrial respiration and the spare respiratory capacity, whereas the Seahorse Real-Time ATP Rate Assay is designed to measure mitochondrial as well as glycolytic ATP production rates in living cells. A Seahorse 96-well cell culture microplate was coated with poly-L-lysine for 10 min before seeding HeLa cells as at density of 1 × 104. The outer rim of the plate was used for background correction, containing only medium without cells. HeLa cells were incubated for 48 h at 37°C, 5 % CO_2_ in cell culture media before the assay was started. Sensor cartridge was hydrated one day prior to sample analysis by sterile H_2_O at 37°C in a CO_2_-free incubator. One to two hours before starting the analysis, water was replaced by 200 μl of pre-warmed Seahorse XF Calibrant per well. All assay compounds (oligomycin, FCCP, rotenone and antimycin A) were solubilized in DMSO to a stock concentration of 5 mM and stored at -20°C. At the day of the assay, compounds were prepared for loading in XF sensor cartridge. Concentrations were chosen according to the Seahorse assay performed. Stock solution of compounds were mixed with assay medium to generate 10 x port concentrations. Compounds were loaded into ports in XF sensor cartridge. XF sensor cartridge plate was placed into the Seahorse XFe analyzer and calibration was performed. Afterwards, calibration plate was replaced by the XF cell culture microplate and measurement was started. Pre-warmed Seahorse DMEM assay media (pH 7.4) was supplied with 10 mM glucose, 2 mM L-glutamine and 1 mM pyruvate. Cells were washed once with Seahorse assay media before 180 μl of Seahorse assay media was added to each well. Cells were incubated at 37°C in a non-CO_2_ incubator for 45-60 min prior to the assay. For the XF Real-Time ATP Rate Assay performed with HeLa cells, Seahorse DMEM assay media was exchanged again before starting the assay. The analysis of Seahorse Assays was performed by Wave Software. Data values were exported to Excel and further analyzed as described in User Manuals. For the Seahorse Real-Time ATP Rate Assay, the Seahorse Report Generator in Excel was used. Statistical Analysis was performed using GraphPad PRISM (GraphPad Software, San Diego California, USA).

### DNA constructs

The wildtype PINK1-V5 construct was described previously (Exner et al., 2007). It was generated by cloning the human PINK1 cDNA into the pcDNA6 vector via NheI and XhoI. For cloning of the kinase dead PINK1-V5 mutant (K219A, D362A, D384A), the construct was modified by point mutations by using the indicated primers. For cloning of the PINK1-Δ77-V5 mutant, the PINK1 sequence 78-581 was cloned into pcDNA6 vector.

Wildtype human HOIP, HOIL-1L, SHARPIN were described previously (Muller-Rischart et al., 2013). HA-HOIP 1-697 was generated using the indicated primers and then incorporated into pcDNA 3.1-N-HA via EcoRI and XbaI. HA-HOIP ΔNZF1, HA-HOIP ΔNZF2, HA-HOIP ΔNZF1+2, HA-HOIP ΔUBA and HA-HOIP ΔPUB were generated by overlap extension PCR (Meschede et al., 2020; van Well et al., 2019). The amplified fragments were generated using the primers indicated in the Materials section.

### Expression and purification of recombinant proteins

Bacterial expression and purification of GST-tagged proteins (C-terminal HOIP, M1-linked tetra-ubiquitin, tcPINK1), Strep-tagged proteins (OTULIN and UBAN domain) and His-tagged proteins (mouse UBE1 and UBE2L3) were performed by standardized methods (Rasool et al., 2018; Stieglitz et al., 2012). The proteins were expressed in *E. coli* BL21 (DE3)-pLys to an OD 600 of 0.6–0.8 before induction with 0.1 mM isopropyl-β-d-thiogalactopyranoside and a change in temperature to 18°C overnight. In the case of C-terminal HOIP, the medium was supplemented with 150 µM ZnCl_2_. Cells were harvested and resuspended in 20 mM Tris-Cl pH 7.5, 150 mM NaCl, 10% glycerol and 1 mM DTT and lysed by a French press. After centrifugation for 40 min at 38000 g, tagged proteins were affinity-purified with a HisTrap HP nickel column (GE Healthcare) or glutathione Sepharose 4B resin (GE Healthcare) or StrepTrap XT (Cytiva). After that, a further step of purification was implemented by size exclusion chromatography (HiLoad 16/600 Superdex 75 pg, Cytiva) or anionic exchange (Hi-Trap Capto Q, Cytiva). In all purification steps, the eluats were collected, and SDS-PAGE was performed to determine the protein purity. The protein concentration was determined using the absorbance at 280 nm and the extinction coefficient of each construct by Nanodrop (Thermo Fisher). Recombinant ubiquitin was purchased from Sigma Aldrich (79586-22-4).

### Quantitative RT-PCR

Total cellular RNA was isolated with the RNeasy Mini Kit (Qiagen) according to manufacturer’s instructions. 1000 ng RNA was reverse-transcribed into cDNA using the iScript™ cDNA Synthesis Kit (Bio-RAD, CA, USA). cDNA was diluted 1:5 in PCR-grade dH_2_O and 1.5 µl of the cDNA-mix was used per 20 µl real-time PCR reaction with 0.25 µM of each primer and 2 x FastStart Essential DNA Green Master Mix (Roche). For each target and reference gene, samples were run in triplicates on a Light Cycler 96 (Roche). On each plate relevant negative controls were run. Melt curves were studied for all assays, with the Tm checked to be within known specifications for each assay. Standard curves were run for each primer pair. Relative quantification of PINK1 mRNA was performed using the 2^-ΔΔ*CT*^ method.

### Immunocytochemistry

SY-SY5Y, HAP1, MEFs and neuronal cells were cultivated on glass coverslips (Laboratory Glassware Marienfeld). 24-72 h after transfection, cells were washed with PBS pH 7.4, fixed for 15 min with 4% paraformaldehyde in PBS pH 7.4 and permeabilized with 0.2% (v/v) Triton X-100 in PBS for 10 min or with 0.5% saponin, 1% BSA in PBS for 45 min at room temperature. After blocking with 1% BSA or 5% goat/donkey serum, cells were stained with primary antibodies at a dilution of 1:100 to 1:1000 in PBS or 0.5% saponin, 1% BSA in PBS at 4°C overnight, washed with PBS and incubated with fluorescent dye-conjugated secondary antibodies AlexaFluor488, 555 or 647 (Thermo Scientific), at a dilution of 1:1000 for 1 h at room temperature. Cells were washed with PBS pH 7.4 three times and subsequently incubated with DAPI (1µg/mL) in H_2_0. Following a final wash in H_2_O, cells were mounted in Fluoromount G (Thermo Fisher Scientific).

### Mitochondrial and nuclear staining in SPLICS^NU-MT^ transfected cells

Cells were fixed with 3.75% paraformaldehyde in PBS for 20 minutes, permeabilized with 0.3% Triton X-100 for 10 min, blocked with 1% bovine gelatin in PBS for 10 min three times and then washed with PBS for 5 min three times. Mitochondria were stained using a 1:100-diluted rabbit anti-TOM20 primary antibody (Santa Cruz). Cells were incubated with the antibody in a wet chamber for 2 h at room temperature, blocked three times with gelatin in PBS for 10 minutes and washed three times with PBS for 5 min. Cells were then incubated with a 1:100-diluted AlexaFluor 594-conjugated anti-rabbit secondary antibody (ThermoFisher) in a wet chamber for 45 min at room temperature, rinsed with gelatin in PBS for three times, then nuclei were stained by incubating the cells with 1µg/ml Hoechst 33342 in PBS for 10 min. Finally, the cells were washed three times with PBS for 5 min and mounted on glass slides before being analyzed by a Zeiss sp5 confocal microscope.

### Assessment of the mitochondrial membrane potential

For ΔΨ_m_ determination, HeLa cells were stained with 7 nM tetramethylrhodamine-ethyl-ester-perchlorate (TMRE). For ratiometric imaging, dual staining was performed with MitoTracker Green FM (MTG), used at a concentration of 100 nM. MTG is a ΔΨ_m_ independent dye. TMRE is rapidly equilibrated and non-toxic for respiration in those low concentration. Cells grown on glass coverslips were stained in fresh medium for 30 min at 37°C with 100 nM MitoTracker^®^ Green FM, washed once with PBS and once with medium and then imaged in fresh media containing 7 nM TMRE and 10 µM verapamil (after another 30 min of incubation). MTG emission was recorded between 500-510 nm. TMRE was added 30 min before the measurements and kept in medium throughout the experiments. During imaging, no MTG was present in the medium. TMRE was obtained from Biomol GmbH and MTG from ThermoFisher Scientific. Determination of ΔΨ was performed by simultaneous TMRE and MTG measurements (Z-stacks, 0.25 nm steps, 7 slices) to enable normalization according to Wikstrom et al. (Wikstrom et al., 2007) with a cLSM (Leica SP8) equipped with a tunable white light laser (470-670 nm), a 63x water objective (N.A. 1.2) and two highly sensitive hybrid GAsP (HyD) detectors. Two technical replicates were conducted. For evaluation, an Otso mask was used to obtain the mitochondria skeleton only. Mean grey values were determined for each cell and the fluorescence emission ratio (TMRE/MTG) calculated. Box-and-whisker plot from mean grey value data points (each data point represents the mean of one cell = one mitochondrial network). Error bars denote 5^th^ to 95^th^ percentile values, boxes represent 25^th^ to 75^th^ percentiles. Vertical lines in boxes represent the median values, whereas square symbols in boxes denote the respective mean values. Minimum and maximum values are denoted by x. Significance levels were determined by a one-way ANOVA: *p ≤ 0.05; **p ≤ 0.01; ***p ≤ 0.001.

### Live-cell imaging of human primary macrophages

Macrophages were plated on at 50000 cells/well in μ-Slide 8-well chambers (80826, Ibidi, Martinsried, Germany) and incubated with 0.5 µM MitoTracker Green (Thermo Fisher Scientific) for 15 min at 37 °C prior to imaging. The nuclei were stained using 2 µM Hoechst (Thermo Fisher Scientific). TNFα was added immediately before staring video recording. Imaging was carried out using confocal microscope (LSM800, Carl Zeiss, Oberkochen, Germany) and LD Plan-Apochromat 40x objective.

### Electron microscopy

Cell grown on ACLAR^®^–Fluoropolymer fil (Plano) were fixed by immersion using 2 % glutaraldehyde in 0.1 M cacodylate buffer at pH 7.4 overnight at 4°C. After postfixation in 1% osmium tetroxide and pre-embedding staining with 1% uranyl acetate, tissue samples were dehydrated and embedded in Agar 100. Thin sections (80 nm) were counterstained using 1% uranylacetat (aq) and examined using a Talos L120C (Thermo Fisher Scientifc).

### Flow cytometry

Flow cytometry was performed on a BD FACSymphony A5 cell analyzer. HeLa FlpIn TRex cells stably expressing mt-mKEIMA and doxycycline-inducible Parkin were seeded in 6-well plates 48 h prior to treatment. Parkin expression was induced for 24 h by 0.25 µg/ml doxycycline. Cells were treated with TNF for the indicated time (25 ng/ml for 30 min or 10 ng/ml for 16 h) and harvested by Trypsin digestion. 10,000 events were preselected for viable, fluorescent, single cells. Dual-fluorescence for 405 nm (mt-mKEIMA pH 7) and 561 nm (mt-mKEIMA pH 4) excitation and 610/20 nm emission was recorded. The ratio between 561 and 405 nm mt-mKEIMA fluorescence is a measure for mitophagic flux 1.

### Mitochondria fractionation

To extract crude mitochondria, cells were washed with ice-cold PBS buffer 2 times and scraped from the dishes. Cells were then collected and centrifuged at 1000 x g for 1 min at 4°C. Cell pellets were resuspended in RSB buffer (10 mM Tris/HCl, 10 mM NaCl, 1.5 mM CaCl_2_, pH 7.4) for 2-3 min on ice to swell, and then centrifuged at 2400 x g for 5 min at 4°C. The pellets were resuspended in 1:1 mixed RSB/MS buffer (420 mM mannitol, 140 mM sucrose, 10 mM Tris/HCl, 5 mM EDTA, pH 7.4 protease inhibitors, phosphatase inhibitors, N-ethylmaleimide) and lysed with a 23 G needle (10 times) followed by a 26 G needle (5 times). For human macrophages, the cell suspension was lysed with a 30 G needle (5 times) in addition). The lysed cell suspension was diluted 1:1 ratio with MS buffer. The lysates were centrifuged at 1500 x g for 15 min at 4°C. The supernatant was collected and centrifuged at 14k x g for 15 min at 4°C to sediment crude mitochondria. The supernatant was centrifuged at 20k x g for 1 h at 4°C to collect purified cytosol. For further purification, the crude mitochondria were resuspended in 36% Opti-prep solution and transferred to a tube, overlayed with 4 ml 30% Opti-prep solution and 4 ml 10% Opti-prep solution. 1 ml diluent (250 mM sucrose, 6 mM EDTA, 120 mM HEPES, pH 7.4) was added on top of the gradient. The Opti-prep gradients were centrifuged at 30k rpm for 4h at 4°C. After centrifugation, the white fluffy layer between 30% and 10% was collected and diluted with M1 buffer (600 mM sucrose, 50 mM Tris/HCl, 10 mM EDTA, pH 8.0), centrifuge at 14k x g for 10 min at 4°C to sediment purified mitochondria.

### Linear ubiquitination assay

Cells were lysed in denaturing lysis buffer (1% SDS in PBS) and heated (95°C for 10 min). Protein extracts were diluted 1:10 with non-denaturing lysis buffer (1% Triton X-100 in PBS) and cleared by centrifugation. To pull down proteins modified by linear ubiquitin chains, 20 µg of the recombinant UBAN domain of NEMO was added, which carries an N-terminal Strep-Tag II. After incubation for 1 h at 4°C, Strep-Tactin beads were added and incubated overnight at 4°C. Beads were collected by centrifugation and washed. Laemmli sample buffer was added, and the samples were boiled for 10 min. Proteins binding to the UBAN domain were separated with SDS-PAGE and analyzed by Western blotting using an anti-M1 ubiquitin antibody.

### Co-immunoprecipitation

Cells were harvested in cold PBS and subsequently lysed in 1% (v/v) Triton X-100 in PBS supplemented with Protease Inhibitor Cocktail (Roche), N-ethylmaleimide and sodium orthovanadate. The lysates were cleared by centrifugation at 20,000 x g and incubated overnight with anti-HA agarose beads at 4°C with gentle rotation. Beads were washed three times with lysis buffer. Immunopurified proteins were eluted by adding Laemmli sample buffer (LSB) and boiling for 10 min.

For endogenous PINK1 immunoprecipitation, crude mitochondria were extracted from cells, and lysed in 1% Triton lysis buffer on ice for 5 min, then centrifuged at 20k x g for 20 min at 4°C. The lysates were incubated with rabbit anti-PINK1 antibody or IgG epitope control overnight at 4°C. The samples were then incubated with pre-washed protein A beads for 3 h. The protein A beads were washed with 1% Triton lysis buffer 3 times before adding to the samples. The beads were washed three times with 1% Triton lysis buffer, finally eluted with 2 x LSB and analyzed by Western blotting.

### Size exclusion chromatography

HEK293T cells expressing PINK1-V5 were lysed in Tris buffer (50 mM Tris pH 7.5, 1 mM MgCl_2_, 1 mM DTT, and complete protease inhibitor) by repeated passing through a syringe needle. After 15 min incubation on ice, lysates were cleared by centrifugation (20,000 x g, 20 min, 4°C). Protein concentration of the samples was measured by the Bradford protein assay and equal amounts of proteins were loaded on the column. Proteins were separated on a Superdex 200 10/300 GL column using an ÄKTA chromatography system (GE Healthcare Life Sciences). 0.5 ml or 1 ml fractions were collected and analyzed by Western blotting using antibodies against HOIP and V5.

### *In vitro* ubiquitination assay

PINK1 was immunoprecipitated from cell lysates by anti-V5 beads. The beads were washed three times with 1% Triton lysis buffer and twice with ubiquitination reaction buffer (50 mM Tris-HCl, 5 mM MgCl_2_, 0.5 mM TCEP), and then incubated with recombinant ubiquitin (30 µM), mouse Ube1 (0.1 µM), UBE2L3 (1 µM), C-terminal HOIP (4 µM) and ATP (2 mM) in the *in vitro* ubiquitination reaction buffer for 90 min at 30°C. Reactions were terminated by adding 5x LSB into the reaction mix at 97°C for 3 min. Samples were analyzed by Western blotting.

### *In vitro* deubiquitination assay

Recombinant M1-linked tetra-ubiquitin (2.5 µg) was incubated with or without 16 µg tcPINK1 or 16 µg BSA as a control for 2 days at 30°C in kinase buffer (500 mM Tris, 100 mM MgCl_2_, 10 nM ATP, 10 mM DTT). For the DUB assay, 0.26 µg recombinant OTULIN was added incubated at 37°C for the indicated time. The DUB reaction was terminated by 5 x LSB and heating (80°C). The samples were analyzed by Western blotting and Phos-tag™ PAGE.

### Cellular deubiquitination assay

PINK1 was immunoprecipitated from cellular lysates and the beads were resuspended in low salt buffer containing recombinant OTULIN (2 mM), and incubated at 25 °C for 1 h. The reaction was terminated by adding 5x LSB and heating (95°C). The samples were analyzed by Western blotting.

### Apoptosis assays

Activated caspase-3 was quantified as described previously (Meschede et al., 2020). In brief, cells plated on glass coverslips were stained with antibodies against activated caspase-3 by indirect immunofluorescence and signals were visualized by fluorescence microscopy by using a Nikon Eclipse E400 microscope.

For the assessment of Bax recruitment, mitochondrial fractions were analyzed by Western blotting using Bax-specific antibodies. For analyzing cytochrome c release, subcellular fractionation was performed by differential centrifugation. The cytosolic and mitochondrial fractions were analyzed by Western blotting using antibodies against cytochrome c.

### Image acquisition

Fluorescence microscopy was performed using a Zeiss ELYRA PS.1 equipped with an LSM880 (Carl Zeiss, Jena) and a 20x, 63x oil or 100x oil immersion objective or a C2+ system (Nikon). Super-resolution images were generated by structured iIlumination microscopy (SIM) using 405, 488 and 561 nm widefield laser illumination. SIM confocal images were processed using the ZEN2.3 software (Carl Zeiss, Jena). For the p65 translocation assay, laser scanning microscopy was performed using the 405, 488 and 561 laser illumination set in individual channels to avoid cross-talk.

### Image analysis

#### Mitochondrial movement

For the analysis of mitochondrial movement, cells were incubated with MitotrackerGreen™ (50 nM) and Hoechst (5 µg/mL) and imaged every 30 s after treatment with TNF (25 ng/mL). Mitochondria and the nucleus were segmented using Imaris 9.8 Surface and Spot modules and mitochondrial spots were classified with respect to the nuclear distance.

#### Cytosolic/nuclear ratio of the p65 signal

HeLa cells were plated on glass coverslips and transfected with control or Miro1-and Miro2-specific siRNAs. 48 h after transfection cells were treated with TNF (25 ng/ml) for 15 min and then fixed with 4% PFA in PBS for 10 min at room temperature. Cells were permeabilized with 0.2% Triton X-100 for 30 min and blocked with 5% donkey or goat serum in 0.2% Triton X-100 in PBS for 1 h at room temperature. Subsequently, the cells were incubated with an anti-p65 and anti-tubulin antibody diluted in blocking buffer overnight at 4°C. After washing, cells were incubated with an AlexaFluor488 and AlexaFluor555 secondary antibody for 1 h at room temperature. Finally, the cells were washed extensively with PBS and then mounted onto glass slides and analyzed by laser scanning microscopy (LSM). For the analysis of the signal intensity ratio between the cytosol and the nucleus, LSM datasets including stainings for the whole cell (tubulin), p65 and the nucleus (DAPI) were exported using Zeiss Zen Blue (2.1). Based on the individual channels, datasets were segmented in a cytoplasmic and nuclear compartment and the fluorescence p65 signal intensity ratio for every cell was measured using CellProfiler 4.2.1.

Alternatively, translocation of p65 from the cytosol to the nucleus was determined by indirect immunofluorescence based on single cell analysis. SH-SY5Y cells or MEFs were plated on glass coverslips and transfected with control or PINK1-specific siRNA. For rescue experiment, cells were co-transfected with WT PINK1, PINK1Δ77, or kinase-dead PINK1-K/D mutant. Two days after transfection, the cells were treated with TNF (25 ng/ml, 25 min) and then fixed with 4% PFA in PBS for 10 min at room temperature. Cells were permeabilized with 0.2% Triton X-100 for 10 min and blocked with 5% donkey or goat serum in 0.2% Triton X-100 in PBS for 1 h at room temperature. Subsequently, the cells were incubated with an anti-p65 antibody diluted in blocking buffer overnight at 4°C. After extensive washing, cells were incubated with an ALEXA488-conjugated secondary antibody for 1 h at room temperature. Finally, the cells were washed extensively with PBS and then mounted onto glass slides and analyzed by fluorescence microscopy using a Nikon Eclipse E400 microscope. Nuclei were counterstained with DAPI (Sigma) and transfected cells were visualized by EGFP, mCherry or mitoDsRed plasmid cotransfection.

#### Mitochondrial ultrastructure

High resolution images of mitochondria were imported into Imaris 9.8 and scale was adjusted using the correct pixel sizes. For the analysis of average cristae number/mitochondrial area, Imaris 9.8 surface function was used, for the average cristae length data was collected using a direct measure function.

### Mass spectrometry-based analysis for the identification OTULIN interaction partners

In-gel digestion was performed as previously described (Sima et al., 2021). 200 ng tryptic peptides were subsequently measured by nano-LC-ESI-MS/MS. An UltiMate 3000 RSLC nano-LC system (Thermo Scientific, Bremen, Germany) was utilized for nano-HPLC analysis using the following solvent system: (A) 0.1% FA; (B) 84% ACN, 0.1% FA. Samples were loaded on a trap column (Thermo, 100 μm × 2 cm, particle size 5 μm, pore size 100 Å, C18) with a flow rate of 30 μl/min with 0.1% TFA. After sample concentration and washing, the trap column was serially connected with an analytical C18 column (Thermo, 75 μm × 50 cm, particle size 2 μm, pore size 100 Å), and the peptides were separated with a flow rate of 400 nl/min using a solvent gradient of 4% to 40% B for 98 min at 60°C. After each sample measurement, 1 h of column washing was performed for equilibration. The HPLC system was online connected to the nano-electrospray ionization source of a LTQ Orbitrap Elite mass spectrometer (Thermo Scientific, Bremen, Germany). The mass spectrometer was operated in a data-dependent mode with the spray voltage set to 1,600 V in positive mode and a capillary temperature of 275°C. Full scan MS spectra (mass range 300-2000 m/z) were acquired in the Orbitrap analyzer at a mass resolution of 60,000. The twenty most intensive ions per spectra were subsequently fragmented using collision-induced dissociation (35% normalized collision energy) and scanned in the linear ion trap. The m/z values triggering MS/MS were set on a dynamic exclusion list for 30 seconds.

Proteins were identified and quantified using MaxQuant (v. 1.6.17.0, https://maxquant.org/) using Andromeda as a search engine. Spectra were matched against UniProt/Swiss-Prot using human taxonomy (released 2021_02). Methionine oxidation was set as variable modifications; cysteine carbamidomethylation as a fixed one. The minimum number of peptides and razor peptides for protein identification was 1; the minimum number of unique peptides was 0. Protein identification was performed at a protein false discovery rate of 0.01. The “match between runs” option was on. Intensity-based absolute quantification (iBAQ) was used to estimate protein abundances (Tyanova et al., 2016). The Gene Ontology Cellular Compartment (GOCC) annotation was performed using DAVID Bioinformatics Resources 6.8 (Huang da et al., 2009a, b). Factory settings were maintained. Resulting tables were exported into excel and the number of proteins from the identified cellular compartments were visualized in pie chart.

### Quantification and statistical analysis

For the quantification of mitochondria-nucleus contact sites, Image analysis and contacts quantification was carried on using the ImageJ distribution Fiji (https://imagej.net/software/fiji/), ImageJ plugin VolumeJ, and custom macros written for interorganelle contacts analysis. The analysis was done as previously described (Cali and Brini, 2021). A ROI was traced around the nucleus of probe-positive cells to quantify only contacts that involve the nuclear envelope. Graphs and statistical analysis of quantification data were performed using GraphPad Prism 8.

Data represent the mean ± SD or SEM from n ≥ 3 biological replicates (as indicated in the figure legends). All statistical analyses were performed by using GraphPad PRISM (Version 5; San Diego, CA, USA). To check the normal distribution of data, the Kolmogorov-Smirnov test was applied. Based on the outcome of the test, appropriate parametric and non-parametric tests were chosen. For the comparison of two independent parametric datasets Student’s t-test was used. For the comparison of more than 2 parametric datasets, one-way ANOVA was applied. To correct for α-error inflation resulting from multiple comparisons, ANOVA was followed by Tukey’s post-hoc Multiple Comparison tests. For the direct comparison of two non-parametric datasets, Wilcoxon-Mann-Whitney (U test) was used. Significance levels for all tests: *p ≤ 0.05; **p ≤ 0.01; ***p ≤ 0.001.

## MATERIALS

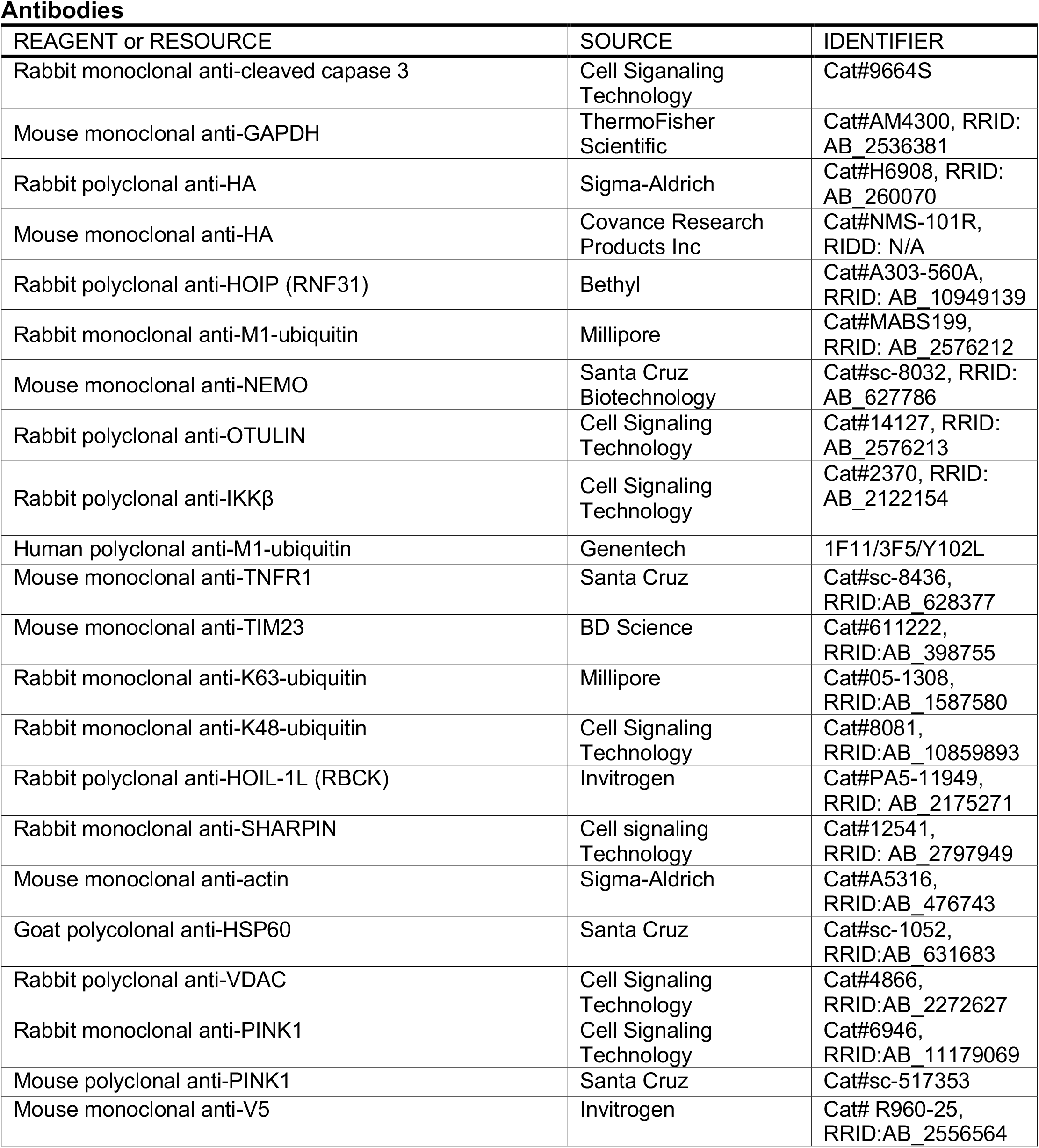
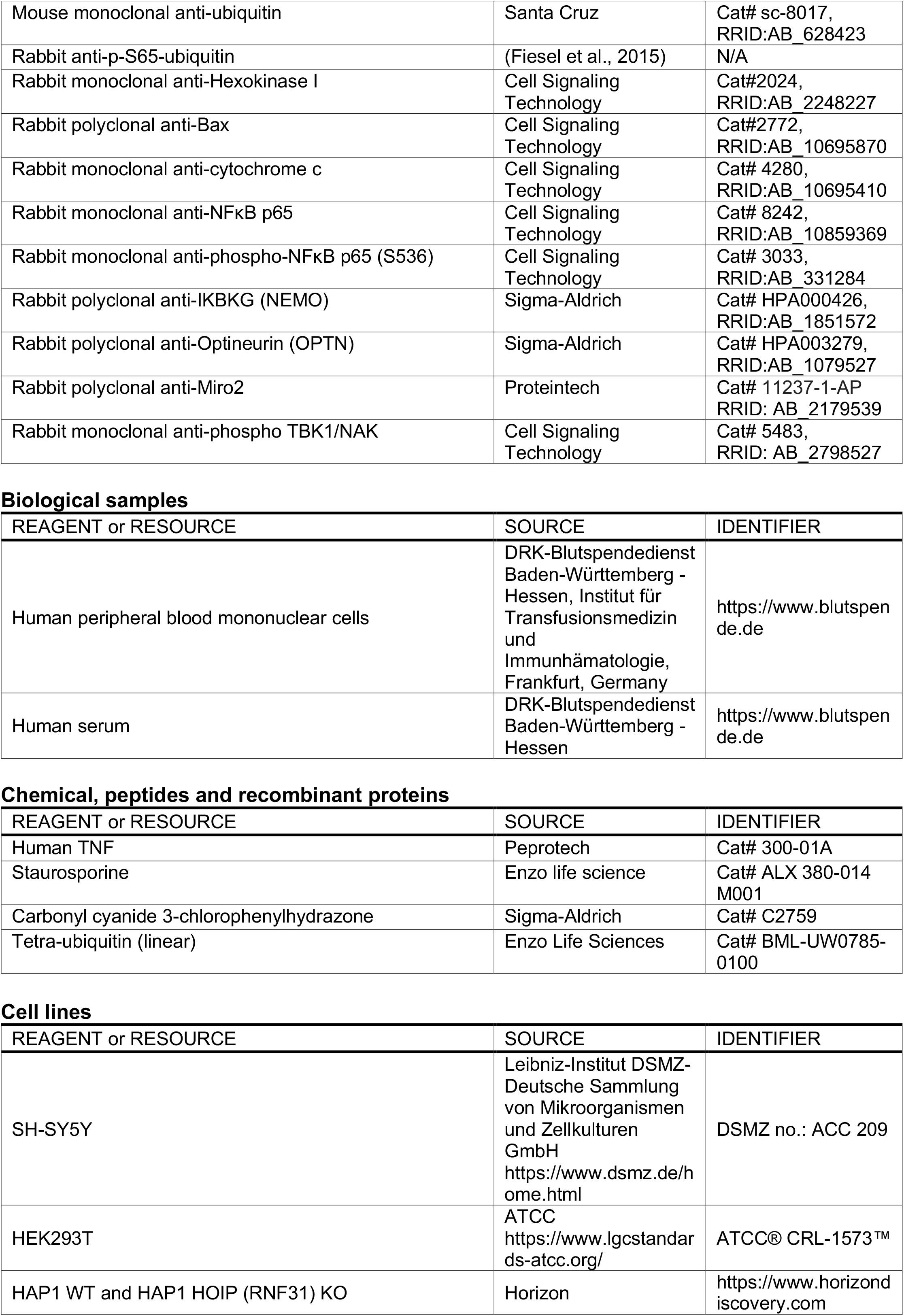
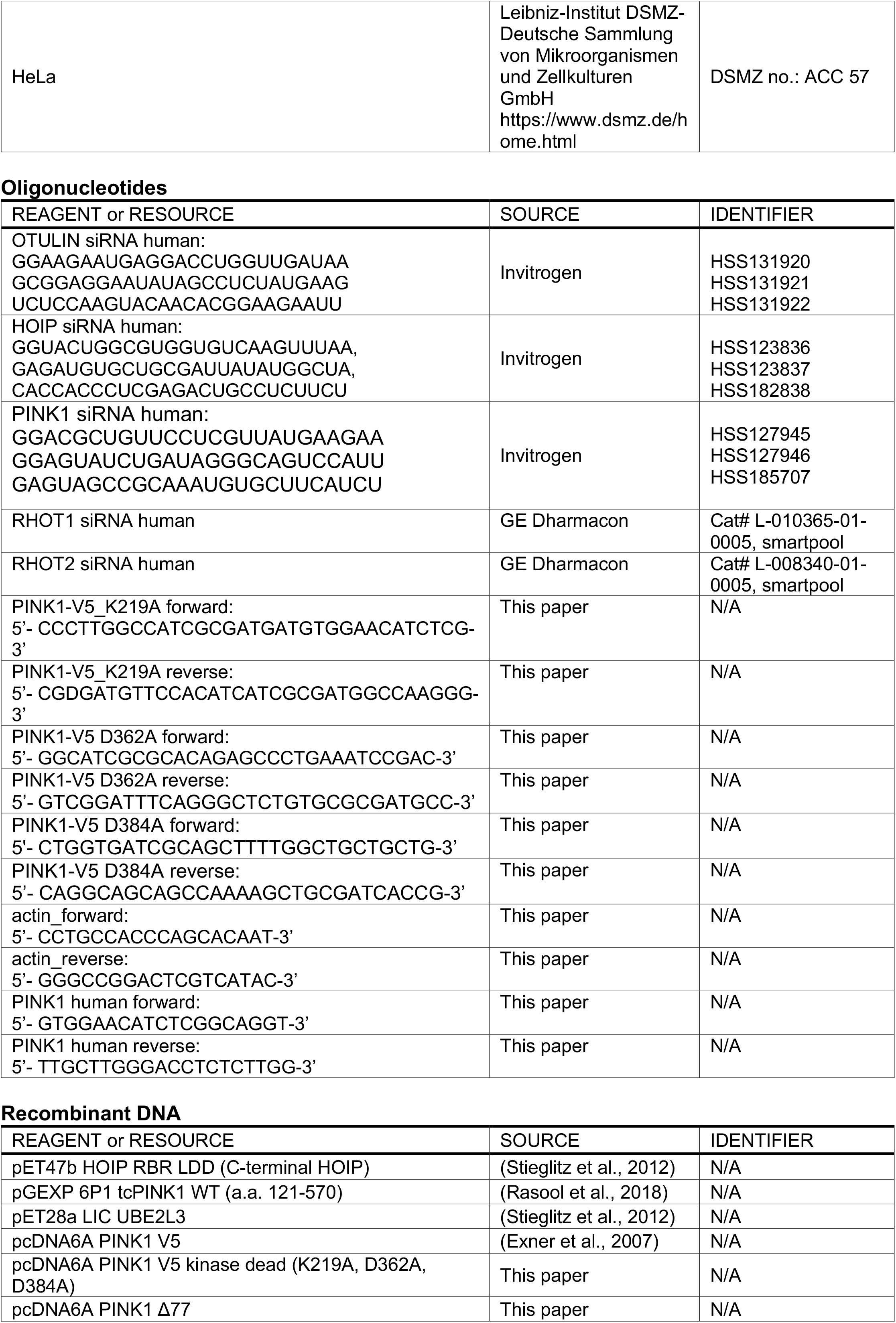
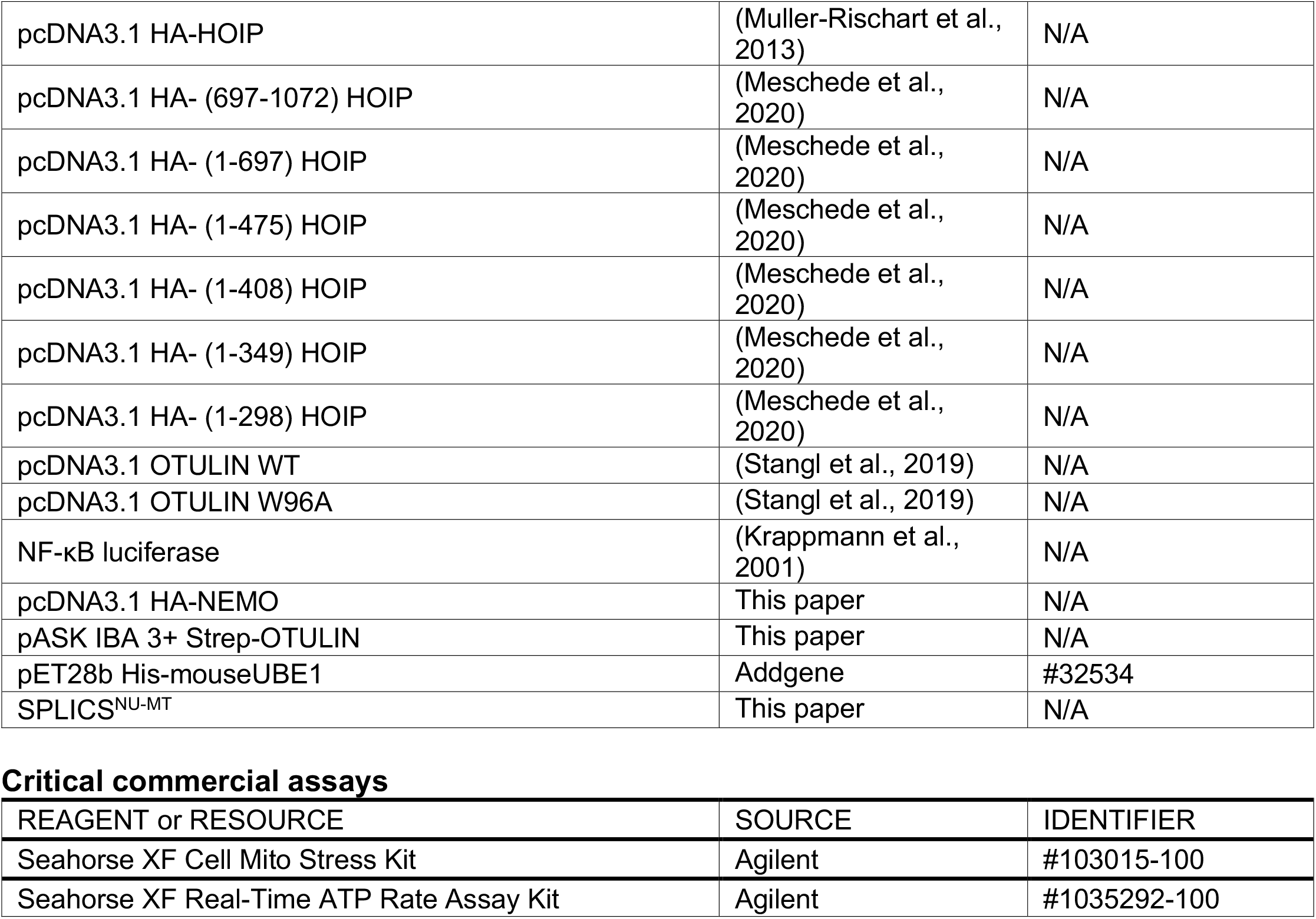

