## Supplementary Figures for "LUBAC assembles a signaling platform at mitochondria for signal amplification and shuttling of NF-ĸB to the nucleus"

**A**

| Accession | Proteine Name | Gene Name | X̄ iBAQ |
|---|---|---|---|
| **Q96BN8** | Ubiquitin thioesterase otulin | OTULIN | 19.52 |
| **Q96L92** | Sorting nexin-27 | SNX27 | 18.75 |
| **O15347** | High mobility group protein B3 | HMGB3 | 16.69 |
| **Q16718*** | NADH dehydrogenase [ubiquinone] 1 alpha subcomplex subunit 5 | NDUFA5 | 15.28 |
| **Q9Y3D3*** | 28S ribosomal protein S16 | MRPS16 | 15.25 |
| **O14828** | Secretory carrier-associated membrane protein 3 | SCAMP3 | 15.23 |
| **O00264*** | Membrane-associated progesterone receptor component 1 | PGRMC1 | 15.21 |
| **P40616** | ADP-ribosylation factor-like protein 1 | ARL1 | 15.19 |
| **P62495** | Eukaryotic peptide chain release factor subunit 1 | ETF1 | 15.14 |
| **P61421** | V-type proton ATPase subunit d 1 | ATP6V0D1 | 14.91 |
| **P20618** | Proteasome subunit beta type-1 | PSMB1 | 14.64 |
| **Q9HB71** | Calcyclin-binding protein | CACYBP | 14.58 |
| **P56556*** | NADH dehydrogenase [ubiquinone] 1 alpha subcomplex subunit 6 | NDUFA6 | 14.51 |
| **P51570** | Galactokinase | GALK1 | 14.44 |
| **P08243** | Asparagine synthetase [glutamine-hydrolyzing] | ASNS | 14.39 |
| **Q9BTV4** | Transmembrane protein 43 | TMEM43 | 14.38 |
| **O15173** | Membrane-associated progesterone receptor component 2 | PGRMC2 | 14.27 |
| **P30040** | Endoplasmic reticulum resident protein 29 | ERP29 | 14.01 |
| **P00387*** | NADH-cytochrome b5 reductase 3 | CYB5R3 | 13.98 |
| **Q9Y3B4** | Splicing factor 3B subunit 6 | SF3B6 | 13.96 |
| **Q96EP0** | E3 ubiquitin-protein ligase RNF31 | RNF31 | 13.89 |
| **Q9HD45** | Transmembrane 9 superfamily member 3 | TM9SF3 | 13.64 |
| **P82664*** | 28S ribosomal protein S10 | MRPS10 | 13.45 |
| **Q9UHI6** | Probable ATP-dependent RNA helicase DDX20 | DDX20 | 13.44 |
| **P50995** | Annexin A11 | ANXA11 | 13.41 |
| **O95470** | Sphingosine-1-phosphate lyase 1 | SGPL1 | 13.21 |
| **P46977** | Dolichyl-diphosphooligosaccharide--protein glycosyltransferase subunit STT3A | STT3A | 13.13 |
| **O00505** | Importin subunit alpha-4 | KPNA3 | 13.09 |
| **P31153** | S-adenosylmethionine synthase isoform type-2 | MAT2A | 13.04 |
| **P28331*** | NADH-ubiquinone oxidoreductase 75 kDa subunit | NDUFS1 | 12.93 |
| **O60841** | Eukaryotic translation initiation factor 5B | EIF5B | 12.91 |
| **O14734** | Acyl-coenzyme A thioesterase 8 | ACOT8 | 12.89 |
| **Q8NF37** | Lysophosphatidylcholine acyltransferase 1 | LPCAT1 | 12.83 |
| **P36507** | Dual specificity mitogen-activated protein kinase kinase 2 | MAP2K2 | 12.79 |
| **Q8IXH7** | Negative elongation factor C/D | NELFCD | 12.75 |
| **P82663*** | 28S ribosomal protein S25 | MRPS25 | 12.66 |
| **Q01581** | Hydroxymethylglutaryl-CoA synthase | HMGCS1 | 12.65 |
| **P48730** | Casein kinase I isoform delta | CSNK1D | 12.63 |
| **Q86W42** | THO complex subunit 6 homolog | THOC6 | 12.62 |
| **Q9NRK6*** | ATP-binding cassette sub-family B member 10 | ABCB10 | 12.50 |
| **Q9H5Q4*** | Dimethyladenosine transferase 2 | TFB2M | 12.49 |
| **P04350** | Tubulin beta-4A chain | TUBB4A | 12.32 |
| **Q96EY7*** | Pentatricopeptide repeat domain-containing protein 3 | PTCD3 | 12.16 |
| **Q96DV4*** | 39S ribosomal protein L38 | MRPL38 | 12.15 |
| **Q9BUQ8** | Probable ATP-dependent RNA helicase DDX23 | DDX23 | 12.11 |
| **Q8N163** | Cell cycle and apoptosis regulator protein 2 | CCAR2 | 12.04 |
| **Q8WWY3** | U4/U6 small nuclear ribonucleoprotein Prp31 | PRPF31 | 11.90 |
| **P23921** | Ribonucleoside-diphosphate reductase large subunit | RRM1 | 11.59 |
| **Q9Y6N5*** | Sulfide:quinone oxidoreductase | SQRDL | 11.52 |
| **Q06210** | Glutamine--fructose-6-phosphate aminotransferase | GFPT1 | 11.49 |
| **Q05397** | Focal adhesion kinase 1 | PTK2 | 11.33 |
| **P25685** | DnaJ homolog subfamily B member 1 | DNAJB1 | 11.26 |
| **Q969Z0*** | FAST kinase domain-containing protein 4 | TBRG4 | 11.21 |
| **Q3SY69*** | Mitochondrial 10-formyltetrahydrofolate dehydrogenase | ALDH1L2 | 11.10 |
| **O60701** | UDP-glucose 6-dehydrogenase | UGDH | 11.09 |
| **O94826*** | Mitochondrial import receptor subunit TOM70 | TOMM70A | 10.89 |
| **Q13085** | Acetyl-CoA carboxylase 1 | ACACA | 10.26 |
| **O75122** | CLIP-associating protein 2 | CLASP2 | 10.06 |
| **Q6P0Q8** | Microtubule-associated serine/threonine-protein kinase 2 | MAST2 | 9.90 |
| **Q29RF7** | Sister chromatid cohesion protein PDS5 homolog A | PDS5A | 9.82 |

**B**

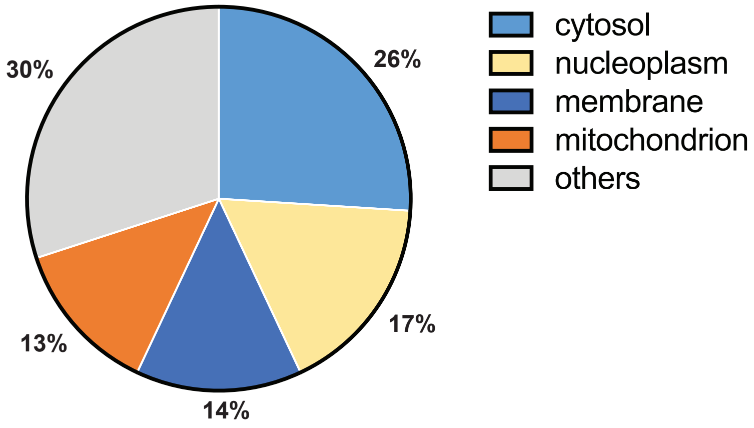

### Suppl. Figure 2

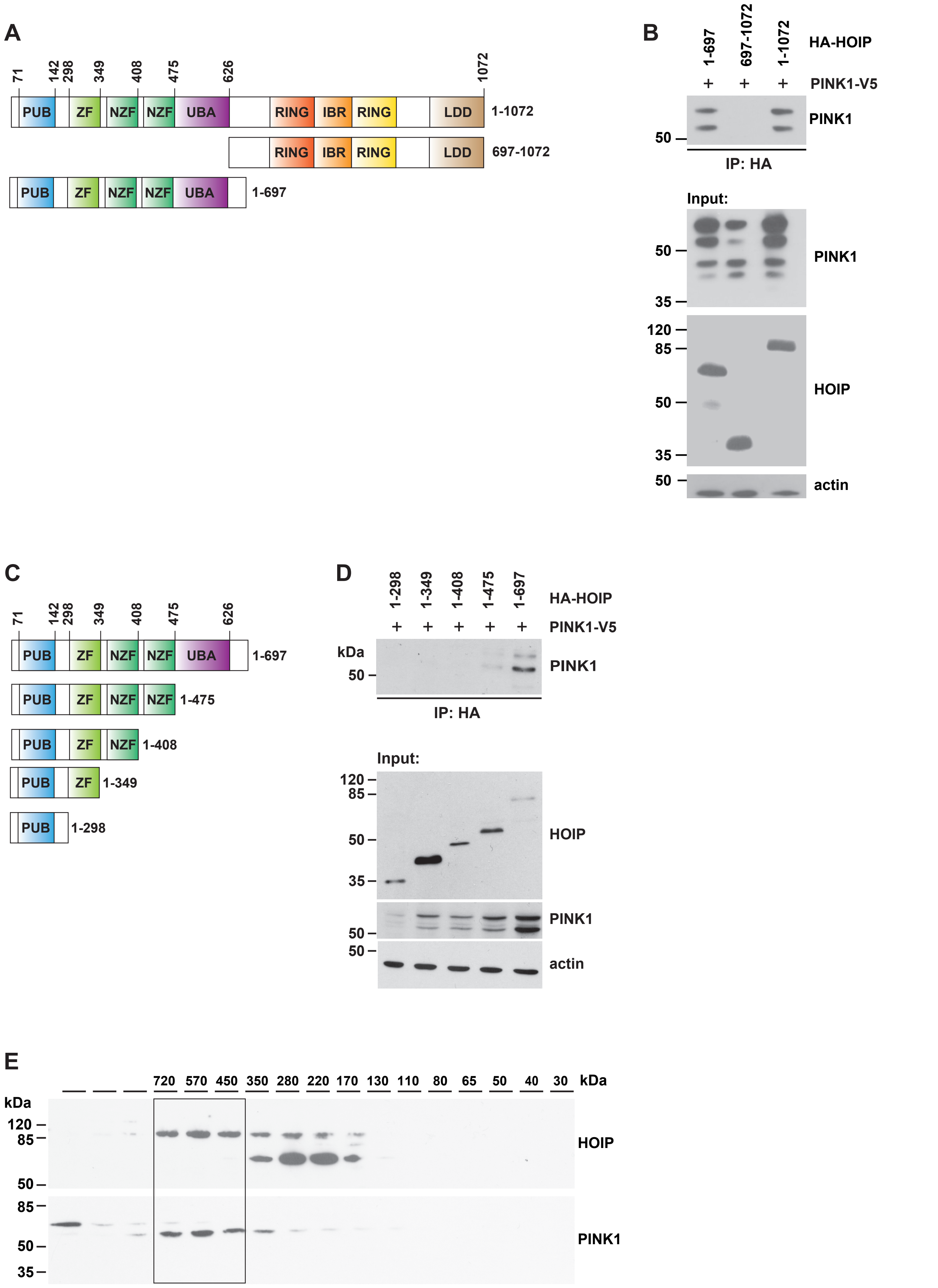

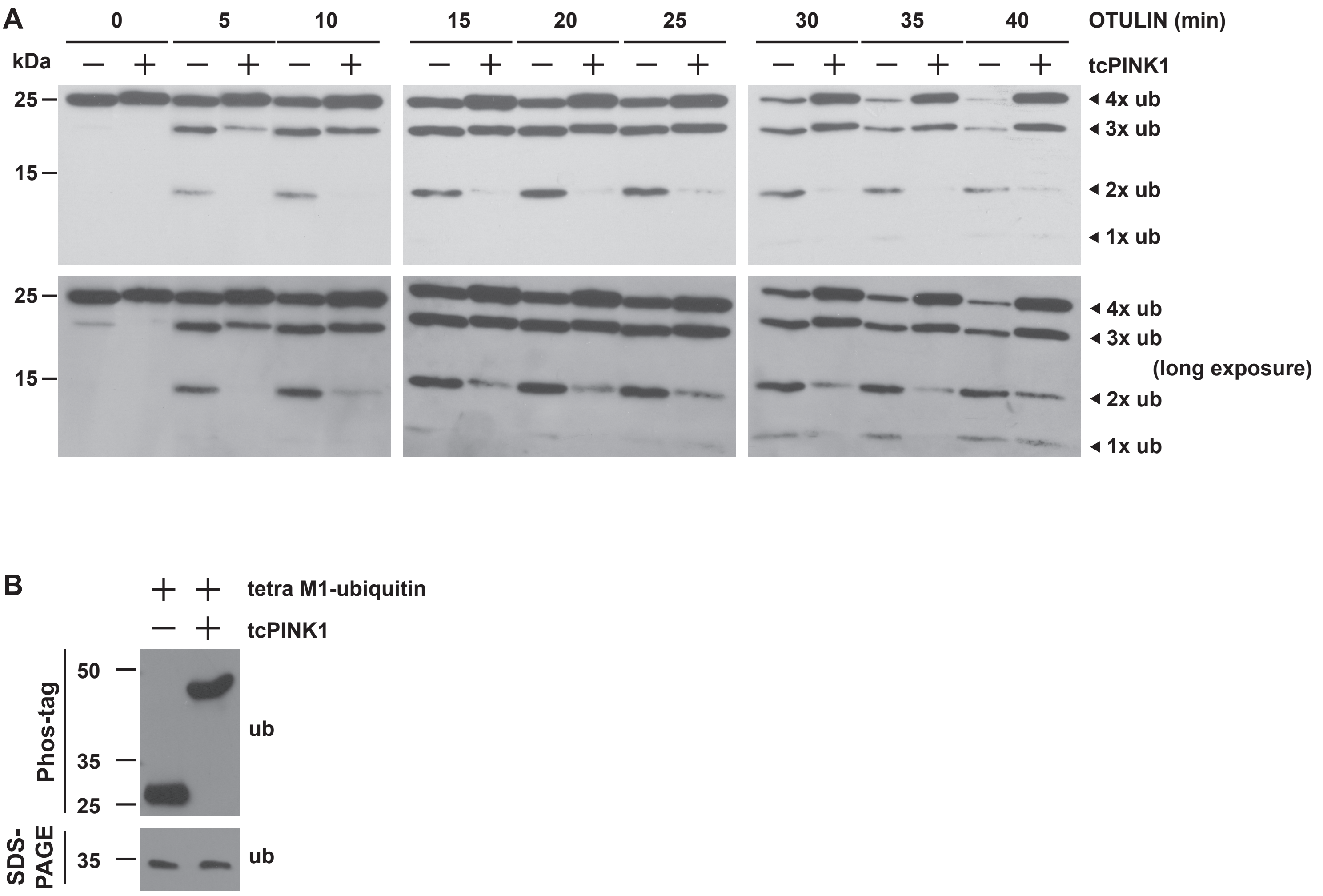
A
0
5
10
15
20
25
30
35
40
OTULIN (min)
kDa
tcPINK1
25
15
4x ub
3x ub
2x ub
1x ub
(long exposure)
B
tetra M1-ubiquitin
tcPINK1
Phos-tag
50
35
25
ub
SDS-PAGE
35
ub

### Suppl. Figure 4

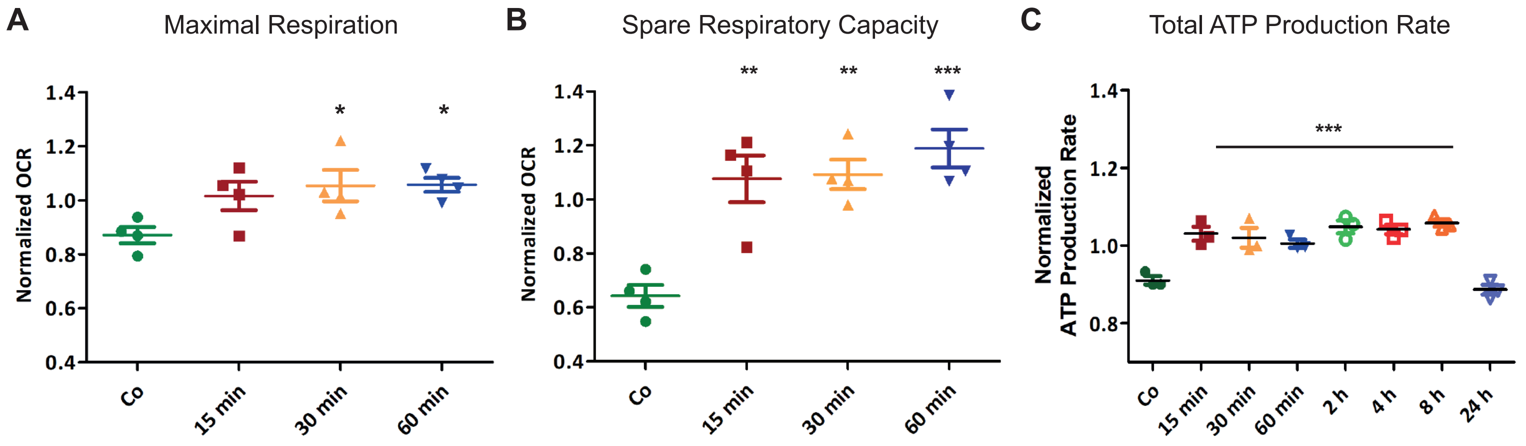

**Suppl. Figure 5**

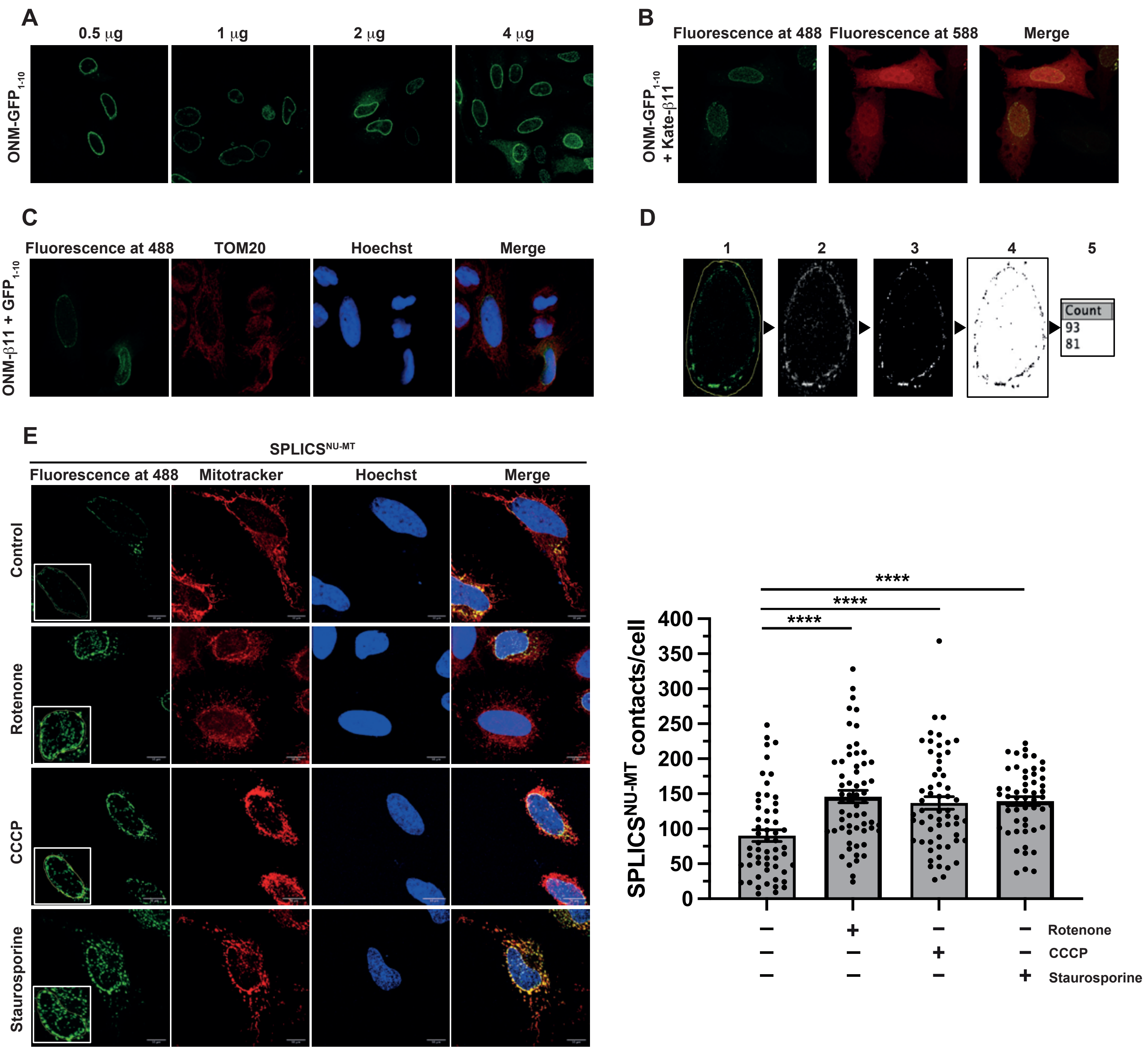

#### SUPPLEMENTARY FIGURE LEGENDS

**Suppl. Figure 1. Identification of OTULIN-interacting proteins by mass spectrometry**

**S1A. OTULIN-interacting proteins.** Proteins identified by co-immunoprecipitation coupled to mass spectrometry analysis using strict filter criteria. Only proteins that were not identified in the control and that were reliably identified in both OTULIN co-immunoprecipitations by at least two unique peptides are listed according to their abundance (iBAQ-value). Mitochondria-associated proteins are marked with asterisk.

**S5A.** Assessment of the nuclear envelope localization of the ONM-targeted $GFP_{1\text{-}10}$ fragment at different concentrations of plasmid transfected in HeLa cells. The $GFP_{1\text{-}10}$ fragment was stained with a mouse anti-GFP primary antibody (Santa Cruz, 1:100) followed by an anti-mouse AlexaFluor 488-conjugated secondary antibody (ThermoFisher, 1:100).

**S5B.** Assessment of the self-complementation capability of the ONM targeted $GFP_{1\text{-}10}$ fragment with a cytosolic mKate-tagged β11.

**S5C.** Nuclear envelope localization and self-complementation of the ONM-targeted β11 fragment co-transfected with an untargeted, cytosolic GFP1-10 fragment. Nuclei were stained with Hoechst33342 (ThermoFisher, 1 μg/ml).

**S5D.** Schematic representation of the contact quantification process using ImageJ plugin VolumeJ and custom-made macros specifically developed for the analysis: 1. a ROI is traced around the nucleus of the cell; 2. the first macro is applied, which clears all the signal outside the ROI and convolves the signal inside; 3. VolumeJ is used to generate a rendering of the convolved signal; 4. the second custom macro is applied on the first and the last frame of the rendering; 5. the contacts per each frame are quantified and the overall number of contacts per cell is calculated by averaging the two results.

**S5E.** Representative images and contact quantification of HeLa cells after induction of the mitochondrial retrograde response (MRR) upon treatment with rotenone (5 μM for 3 h), CCCP (10 μM for 3 h) or staurosporine (1 μM for 2 h). Mitochondria were stained with MitoTracker Red CMXRos (ThermoFisher, 100 nM) in HBSS for 30 minutes. Nuclei were stained as described in C.

**Movie 1. TNF induces the movement of peripheral mitochondria towards the nucleus.** SH-SY5Y cells were stained with NucSpot Live 488 and MitoView 650 for 30 min. After TNF treatment (25 ng/ml) the cells were imaged every 2 min for 30 min. Image analysis was performed by Imaris 9.8 using the Surface (nucleus) and Spot (mitochondria) algorithm. Mitochondrial distance with respect to the nuclear surface was color-coded (purple/blue = proximal; red/yellow =distal).
